# Causal identification of single-cell experimental perturbation effects with CINEMA-OT

**DOI:** 10.1101/2022.07.31.502173

**Authors:** Mingze Dong, Bao Wang, Jessica Wei, Antonio H. de O. Fonseca, Curt Perry, Alexander Frey, Feriel Ouerghi, Ellen F. Foxman, Jeffrey J. Ishizuka, Rahul M. Dhodapkar, David van Dijk

## Abstract

Recent advancements in single-cell technologies allow characterization of experimental perturbations at single-cell resolution. While methods have been developed to analyze such experiments, the application of a strict causal framework has not yet been explored for the inference of treatment effects at the single-cell level. In this work, we present a causal inference based approach to single-cell perturbation analysis, termed CINEMA-OT (Causal INdependent Effect Module Attribution + Optimal Transport). CINEMA-OT separates confounding sources of variation from perturbation effects to obtain an optimal transport matching that reflects counterfactual cell pairs. These cell pairs represent causal perturbation responses permitting a number of novel analyses, such as individual treatment effect analysis, response clustering, attribution analysis, and synergy analysis. We benchmark CINEMA-OT on an array of treatment effect estimation tasks for several simulated and real datasets and show that it outperforms other single-cell perturbation analysis methods. Finally, we perform CINEMA-OT analysis of two newly-generated datasets: (1) rhinovirus and cigarette smoke-exposed airway organoids, and (2) combinatorial cytokine stimulation of immune cells. In these experiments, CINEMA-OT reveals potential mechanisms by which cigarette smoke exposure dulls the airway antiviral response, as well as the logic that governs chemokine secretion and peripheral immune cell recruitment.

## 1 Introduction

Cellular responses to environmental signals are a fundamental component of biological functioning, playing an integral role in both homeostasis and disease [1]. For decades, controlled perturbation experiments have been used to reveal the underlying mechanisms of biological processes. Recent advances in single-cell technologies enable complex experiments measuring high dimensional phenotypes at high throughput under diverse stimulation conditions [2–8]. However, deriving biological insights from these experiments remains a challenge.

For the analysis of single-cell perturbation data, several approaches have been developed, which can be categorized as follows:

1. *Differential expression* methods test for statistically significant differences in gene expression between cell populations. [9–13]
2. *Differential abundance* methods perform differential abundance testing in a continuous cellular state manifold. [14–17]
3. *Perturbation analysis* methods aim to reveal underlying biological processes by modeling perturbations in a latent space with either linear models or neural networks. [3–7, 18–21]

While techniques to characterize the effects of perturbations by averaging over populations are routinely used in the analysis of single-cell data, methods allowing for causal single-cell perturbation analysis have not yet been extensively explored. In causal inference, the quantification of responses to perturbations is known as the *treatment effect estimation problem* [22]. In subsequent portions of this manuscript, we will borrow from the terminology of causal inference, referring to perturbations and treatments, as well as response and treatment effect, interchangeably. Ideal causal methods allow for the direct characterization of underlying confounding variation, a feature not provided by existing single-cell analysis tools.

A great deal of variability in cellular responses to treatment may be attributable to underlying confounding variation. [23]. In the case of scRNA-seq experiments, sources of variation such as cell cycle stage, microenvironment, and pre-treatment chromatin accessibility may all act as confounding factors when performing treatment effect estimation [18]. Collectively, confounding factors can be thought of as a cell’s underlying state that may both influence a cell’s gene expression profile, and condition treatment-induced gene signatures. Correct identification of confounders allows for appropriate causal matching of cell pairs between conditions, enabling treatment effect estimation at the single-cell level.

One well-established confounding factor that may affect treatment response is cell type. For example, widely used nucleoside analog chemotherapeutics such as 5-fluorouracil (5-FU) act selectively on cells in the DNA synthesis phase of the cell cycle, killing cancer cells while minimizing effects on healthy tissue [24]. Some mutations may also drive differential response to a stimulation, as is seen with some tumors in response to TGF-β [25]. Confounders may be latent or unobserved, such as different exposures of cells to a drug, which may have different effects at different concentrations within each individual cell.

We aim to solve this problem by introducing a causal framework permitting characterization of perturbation effects at the single cell level. We explicitly model the diversity of cellular responses to perturbation across confounding factors, which does not require prior knowledge of cell identities. With this approach, we not only identify the spectrum of possible cellular responses to a perturbation, but also the underlying confounder state that defines the response pattern for each individual cell.

In this paper, we present CINEMA-OT (Causal INdependent Effect Module Attribution + Optimal Transport), which applies independent component analysis (ICA) and filtering based on a functional dependence statistic to identify and separate confounding factors and treatment-associated factors. CINEMA-OT then applies weighted optimal transport (OT) to achieve causal matching of individual cell pairs. The algorithm is based on a causal inference framework for modeling confounding signals and conditional perturbation effects at the single-cell level. The computed causal cell matching enables a multitude of novel downstream analyses, including but not limited to: individual treatment effect estimation, treatment synergy analysis, sub-cluster level biological process enrichment analysis, and attribution of perturbation effects.

We demonstrate the power of CINEMA-OT by benchmarking it on several simulated and real datasets and comparing it to existing single-cell level perturbation analysis methods. We then perform CINEMA-OT analyses of two newly-generated datasets. In the first, we examine the effects of viral infection and cigarette smoke on innate immune responses in airway organoids. In the second, we perform combinatorial cytokine stimulation of *ex vivo* peripheral blood mononuclear cells in order to characterize how cytokines act in concert to shape immune responses.

## 2 Results

### 2.1 Confounder signal matching via CINEMA-OT

To perform causal single-cell perturbation effect inference, we have adopted the *potential outcome causal framework* [22, 26]. Ideally, to generate causal assertions about the effect of a perturbation on the transcriptional state of a given cell, we would like to measure the same cell both before and after a perturbation is applied. However, the process of obtaining transcript measurements from single cells is destructive, and an individual cell may only be measured once. A solution is to infer *counterfactual* cell pairs, which are inferred causally-linked pairs—predictions of what a cell in one condition would look like in another condition. The potential outcome framework formalizes this concept by establishing a statistical model that describes outcome variables as a function of confounding factors and treatment-associated factors. In our setting, the inference of single-cell treatment effects means distinguishing the effects of biological variation and treatment on treatment-associated gene expression.

In the potential outcome framework, a key difficulty for general unsupervised causal inference is “the mixing of confounders with outcomes”. In the field of causal inference, this is described as learning with both interventions and latent confounding [27]. In our case, a gene can contribute to confounding variation as well as treatment-associated variation. To apply the tools of classical causal inference, confounding factors must first be distinguished from treatment-associated factors. Notably, confounding factors and treatment-associated factors may be treated as a low dimensional function of the gene space.

To unmix confounding effects and treatment-associated effects, we propose two sufficient assumptions, which can be potentially relaxed:

**Assumption 1: Independent sources and noise.** *Confounding factors and treatment events are pairwise independent random variables*. This assumption relies on treated and untreated cells being drawn from the same underlying set of cells. In practice, this is a central part of most single-cell experimental designs.
**Assumption 2 (Informal): Linearity of source signal combinations.** *The expression of each gene can be linearly decomposed as the sum of confounding signal contributions and treatment-associated signal contributions*. In assumption 2, our key restriction on data is that the expression of each gene is a linear combination of underlying signals, which serves as a common ground for most linear data analysis approaches (such as PCA). While this assumption is mainly developed for ensuring the ICA identifiability, we note that it is reasonable to assume that the gene expression of a cell is driven by a mixing of underlying independent biological processes, as considered by various previous works [4, 28, 29]. In addition, we implicitly assume the underlying model’s dimensionality is correctly specified after preprocessing. While this cannot hold precisely for real data, a rank selection procedure can be performed in order to maximize the consistency between the number of PC dimensions and the ground truth dimensionality (See Methods).

Based on these assumptions, we are able to provide the theoretical foundation that the confounding factors of data through ICA are identifiable with a statistical test on each component (see supplementary notes). In CINEMA-OT, a Chatterjee’s coefficient-based distribution-free test is used to quantify if each component correlates with the treatment event [30] (Figure 2A).

Finally, using the identified confounding factors, we apply optimal transport with entropic regularization to generate causally matched counterfactual cell pairs. This is equivalent to applying optimal transport on the full ICA embedding while setting the treatment-associated factors to be zero. Optimal transport is a natural choice for this matching procedure as it preserves mass, is robust to outliers, and avoids collapsing matches at the boundaries of separated clusters within the data [31, 32]. While solving the optimal transport problem is often prohibitively resource-intensive for large-scale biological data, CINEMA-OT considers the tractable case of entropic regularization [33, 31]. Optimal transport with entropic regularization can be formulated as a convex optimization problem which can be solved efficiently using the alternating direction method (Sinkhorn-Knopp algorithm [33, 31]).

### 2.2 Causal matching in the setting of differential abundance

A treatment may change the distribution of cell densities, e.g. cells may die or proliferate in response to some perturbation. That is, there may be differential confounder abundance across experimentally perturbed datasets. Differential abundance can affect the performance of CINEMA-OT since in this case the underlying confounders are no longer independent of the treatment event and assumption (1) is violated. Our experiments have shown that while CINEMA-OT can tolerate moderate levels of differential abundance, it can fail when high levels of differential abundance are present (Supplementary Figure 5).

To address the issue of differential abundance, we have developed a reweighting procedure called CINEMA-OT-W. In this procedure, before applying ICA, we first align the treated cells by their k nearest neighbors (kNN) in the untreated condition, similar to the Mixscape approach [18]. Although the resulting aligned cell populations may be imperfectly mixed, the kNN alignment process groups together cells with similar confounder characteristics. We then cluster the aligned cells based on the confounder space and subsample them to ensure an equal ratio of treated and untreated cells in each cluster. This reweighting step effectively removes the confounding signal from the treatment event, allowing subsequent application of CINEMA-OT to successfully identify the confounders. CINEMA-OT-W greatly extends the power of the original CINEMA-OT in samples with significant differential abundance across experimental conditions. Notably, it can model the case where counterparts of a significant (sub)population of the unperturbed cells are not present in the perturbed data, enabling causal effect identification for datasets with significant differential abundance.

We note that this functionality should be used only when required. When dealing with data exhibiting differential abundance, the ICA’s identifiability is no longer guaranteed, meaning any existing model, including CINEMA-OT-W, may have reduced ability to accurately identify certain classes of cellular responses. For instance, CINEMA-OT-W may preclude the possibility of a cell type converting to another type in response to treatment, in which case the cell type is a treatment-induced factor rather than a confounding factor. Additionally, selecting the optimal resolution of clustering in CINEMA-OT-W may require prior biological knowledge, since suboptimal choices of clustering resolution may result in reduced power to identify distinct cell populations. As an alternative to CINEMA-OT-W, CINEMA-OT also provides an option to assign weights according to user-provided labels (e.g. cell-types). In this case, CINEMA-OT can sample data using confounder labels instead of automatically balancing over all possible covariates.

### 2.3 Causal matching enables various downstream analyses

The matched counterfactual cell pairs computed by CINEMA-OT define two key outputs: (1) the matching correspondence matrix across treatment conditions, and (2) the individual treatment effect (ITE) for each cell with its counterfactual pair across treatments (Figure 1A).

**Figure 1:**
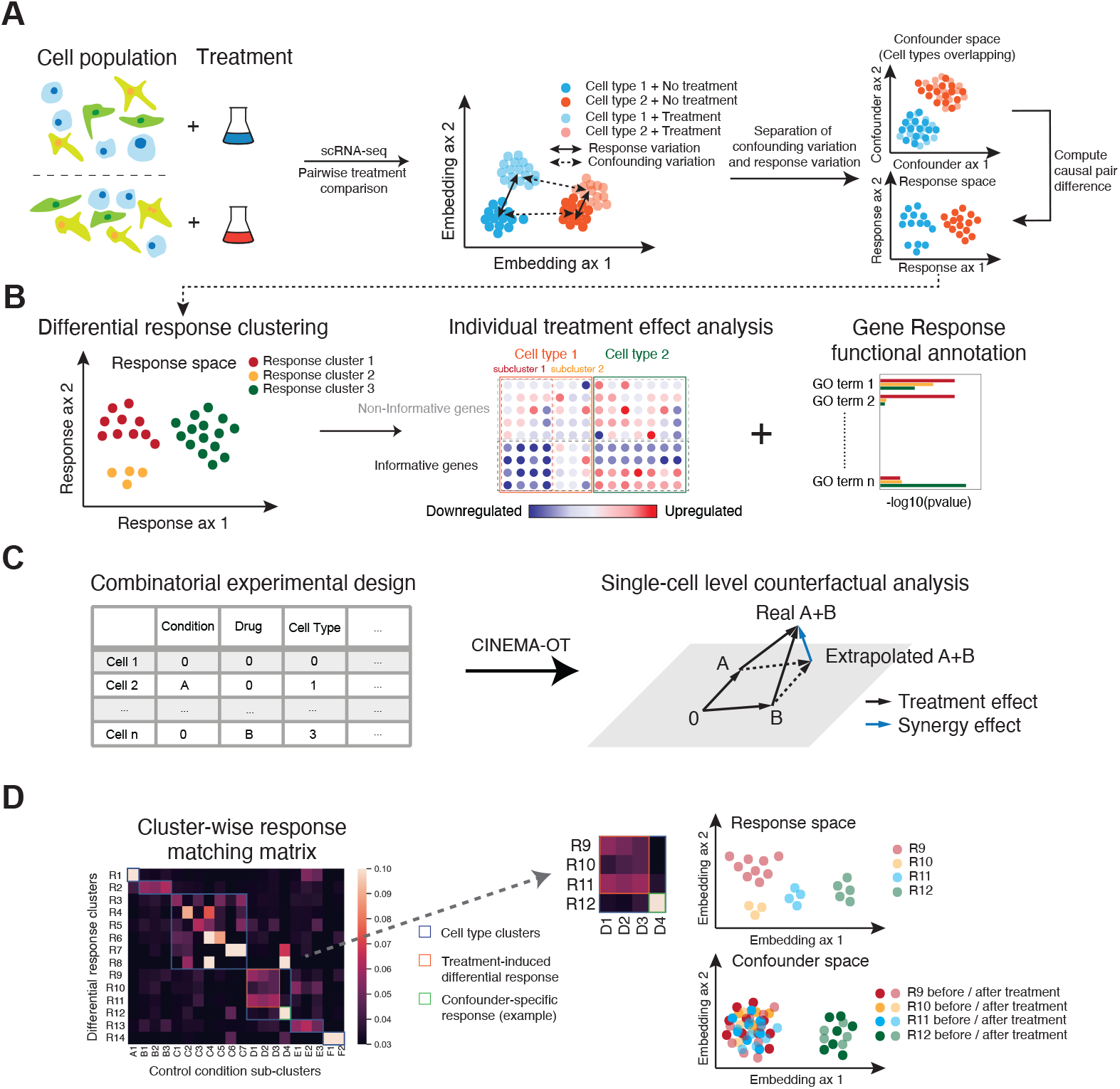
A causal framework for single-cell level perturbation effect analysis. **A.** In single-cell experiments, cells are separated by both treatment batches and latent cell states. Latent states of cells have confounding effects on the response to perturbation. Upon successfully separating confounding variation and response variation in the measurements, we can identify confounding signals for each cell and match counterfactual cell pairs across conditions to compute causal perturbation effects, which indicates the response for each cell. **B.** After characterizing the response matrices at a single-cell level, we can subcluster cells by treatment responses. These responses may be further characterized by other tools, such as gene set enrichment analysis. **C.** We are able to quantify synergy in combinatorial perturbations by evaluating the dissimilarity of extrapolated phenotypes and true combinatorially perturbed phenotypes. **D.** CINEMA-OT can attribute divergent treatment effects to either explicit confounders or latent confounders by analysis of cluster-wise response matching matrices.

**Figure 2:**
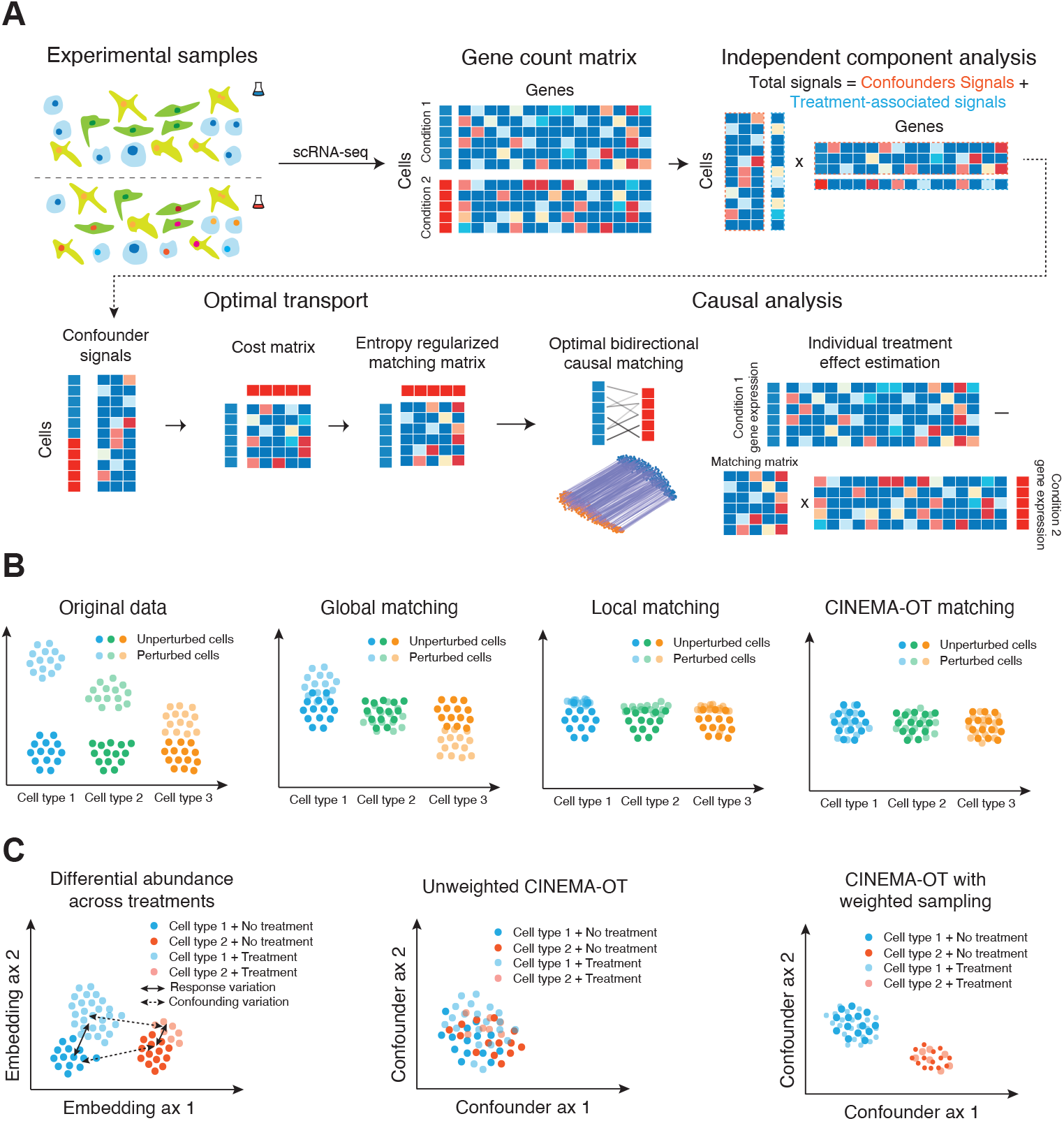
Overview of the CINEMA-OT framework. **A.** scRNA-seq count data is first decomposed into confounder variation and treatment-associated variation using ICA. Cells are then matched across treatment conditions by entropy-regularized optimal transport in the confounder space to generate a causal matching plan. The smooth matching map can then be used to estimate individual treatment effects. **B.** Illustration of the properties of CINEMA-OT compared to other common matching schemes. Global matching may have poor performance when there are confounder-specific heterogeneous responses to treatment. Local matching may be susceptible to boundary effects. By contrast, CINEMA-OT balances these concerns by enforcing preservation of distributional mass. **C.** Differential abundance effects may cause spurious matching by CINEMA-OT. When such effects are present, iterative reweighting may be used to balance cell populations and learn true underlying confounding signals.

Individual treatment effect matrices are cell by gene matrices which can be clustered and visualized by existing scRNA-seq computational pipelines. By clustering over an ITE matrix, we can identify groups of cells with shared treatment response. We may perform statistical analysis to identify the genes with significant magnitudes in response for each group and identify their coordinated biological function by gene set enrichment analysis (Figure 1B).

In addition, when experimental data are available for multiple treatments performed in combination (e.g. control, treatment A, treatment B, combined treatment A+B), we are able to define a synergy effect metric by comparing the predicted effect of combining multiple treatments with the observed effect of combined treatment (Figure 1C). We define this synergy metric by estimating the difference between the true sample under combined treatment (A+B) and the predicted sample by adding the effects of treatment A and treatment B, thus assuming purely linear, non-interactive effects. If no difference is measured, we may conclude that there are no nonlinear or interaction effects between the treatments. If non-zero synergy is present, this points to some interaction between treatments A and B. Synergy is computed for every cell-gene pair, resulting in a matrix of equivalent form to expression or ITE matrices - a unique feature of CINEMA-OT. Notably, as the synergy serves as a summary statistic of the combinatorial cellular responses, the same synergy value may correspond to a number of different underlying mechanisms. For instance, synergistic activation (*x_A+B_* > *x_A_* = *x_B_* = *x_control_* = 0) and uniform inhibition (*x_control_* > *x_A+B_* ≈ *x_A_* ≈ *x_B_*) may lead to the same level of positive synergy for gene x. Our synergy metric enables unbiased investigations of non-linear treatment effects.

Another important task in perturbation effect analysis is the attribution of treatment effects. Differential response can be driven either by differences in explicit confounding factors or by latent factors, such as treatment heterogeneity. Because CINEMA-OT provides a single-cell level matching as one output, the task can be solved by analysis on the clustered matching matrix. Responses that cluster both in response as well as confounder space may be attributed to explicit confounding factors. Conversely, responses that cluster well in the response space but do not demonstrate clustering in the confounder space may be attributed to latent factors (Figure 1D). Such an analysis can be performed either at the cell type level or at the sub-cluster level to reveal underlying heterogeneity. In order to further identify genes with significant explicit confounder-specific treatment effects, we quantify the confounder effect size via a causal regression model and estimate significance using the ratio of confounder-explained effect size to the residual norm (see Methods for additional details).

### 2.4 Validation of CINEMA-OT using simulated data

There are a number of existing methods that perform perturbation effect analysis in single-cell omics data. Several pioneering works in the field propose the use of factor models to identify pseudo-bulk level perturbation effects, including LRICA [4], MIMOSCA [3], scMAGeCK [5], MUSIC [6], and WGCNA/STM [7]. More recently, Mixscape [18] estimates single-cell level perturbation effects by matching between neighboring cells in the shared k-NN graph across conditions. Additionally, several deep learning-based frameworks learn perturbation responses from scRNA-seq data via autoencoders as deep factor models, including scGen [34], compositional autoencoder (CPA) [19], and ContrastiveVI [35]. Finally, CellOT [20] proposes a neural network that learns a non-linear transport map aligning cells from different treatment conditions.

Despite this, none of the existing methods achieve guaranteed confounder identification, which provides interpretable causal effect estimation by aligning cells with the same confounder states across conditions. Among the alternative single-cell treatment effect analysis methods, Mixscape and CellOT do not model the confounding variation. Moreover, Mixscape considers the nearest neighbor relationship on the entire gene expression space instead of distributional matching, which may lead to vulnerability to cell outliers and unbalanced mixing. While the auto-encoder based methods model confounder variation in general, they can suffer from the fundamental limitation of un-identifiability [36] (See Supplementary notes for a detailed explanation), which can reduce their power in identifying ground truth confounding variations. CINEMA-OT is the only method that achieves confounder identification as well as distributional matching for the task of single-cell treatment effect analysis.

Independent component analysis (ICA) has found widespread use in the field of causal inference. One of the most established methods among these is LiNGAM [37], which infers the directed causal relationship between features, with the directions derived by combining the independent noise assumption and ICA identifiability. The LiNGAM framework has been applied to a number of tasks, including: causal discovery for time-series data [38, 39], identifying the features most responsible for an intervention [40], and causal learning across multiple groups [41]. The key distinction between these methods and CINEMA-OT is that the LiNGAM-based methods solve a causal discovery task at the feature (gene) level, while CINEMA-OT seeks to identify the causal effect of an intervention on individual observations (cells). Therefore, LiNGAM-based methods are not appropriate for our task but may find applications in gene regulatory network inference [42–44] or the causal discovery of spatially-regulated gene networks in spatial omics data [45–47].

To investigate how CINEMA-OT differs from existing single-cell level perturbation analysis methods in practice, we perform extensive benchmarking on a number of tasks in simulated scRNA-seq data. Our study involves a meticulous comparison of existing methods with CINEMA-OT with or without sampling (CINEMA-OT/CINEMA-OT-W), including Mixscape, scGen, CPA, ContrastiveVI, and CellOT. Moreover, we explore the potential benefits of integrating batch effect analysis into Mixscape analysis, a method we refer to as Harmony-Mixscape. Additionally, we include a direct optimal transport (Full OT), applied on the original data (without separation of treatment-associated and confounding factors) for the purpose of ablation study. Our comparison is based on three categories of metrics:

1. *Cell distribution equalization after treatment effect removal*. In data sets with or without ground truth, we can measure the validity of treatment effects by examining cell distributions in the gene expression space after treatment effect removal. If different treatments are applied to the same confounder distribution (Assumption 1, see Section 2.1), then these distributions should overlap well after treatment effects are removed. Metrics for evaluating treatment effect removal include average silhouette width (ASW) and principal components regression score (PCR).
2. *Differential response cluster preservation*. If a cell population has divergent responses to a perturbation, the cell population would form clustering structures in the response space. Therefore, preservation of such clustering structures in the estimated treatment effects is essential for perturbation effect identification. In this study, we evaluate the cluster preservation level via an adjusted Rand index (ARI) in individual treatment effect (ITE) matrices.
3. *Attribution accuracy*. Differential response patterns can be attributed to either confounder-specific effects (e.g. cell-type specific effects) or latent-factor driven effects (e.g. treatment drug dose distribution). In simulated data, the attribution accuracy can be measured via independence between confounding factors and responses conditioned on ground truth response labels. In our study this is evaluated by the PC regression score (PCR) in ITE matrices.

To obtain data with ground truth, we simulate data using the Python package Scsim [48], which is a python implementation of the Splatter framework [49] with additional support for simulating gene regulation programs as trajectories. The genes in our simulated data are separated into three subsets, corresponding to the underlying trajectory, cell types, and treatment-associated genes respectively.

We model the dependence between confounders and ground-truth treatment effects in two settings: *1. Confounder-specific treatment effects* modeling diverging responses driven by underlying confounders, such as cell-type specific response. *2. Latent-factor driven treatment effects* modeling the differential treatment effect caused by unobservable latent confounders. In our simulated data, the two settings are covered together by modeling the differential response probabilities as conditional distributions on confounder clusters (Figure 3B). Additionally, we have examined the impact of differential abundance on the performance of various methods by selectively subsampling cells from half of the confounder clusters in the treated condition. We refer to this subsampling ratio as the differential abundance (DA) ratio in the following sections. Furthermore, we have investigated the relationship between the performance of single-cell level treatment analysis and the signal-to-noise ratio of a scRNA-seq dataset by downsampling the gene counts of simulated datasets at different levels.

**Figure 3:**
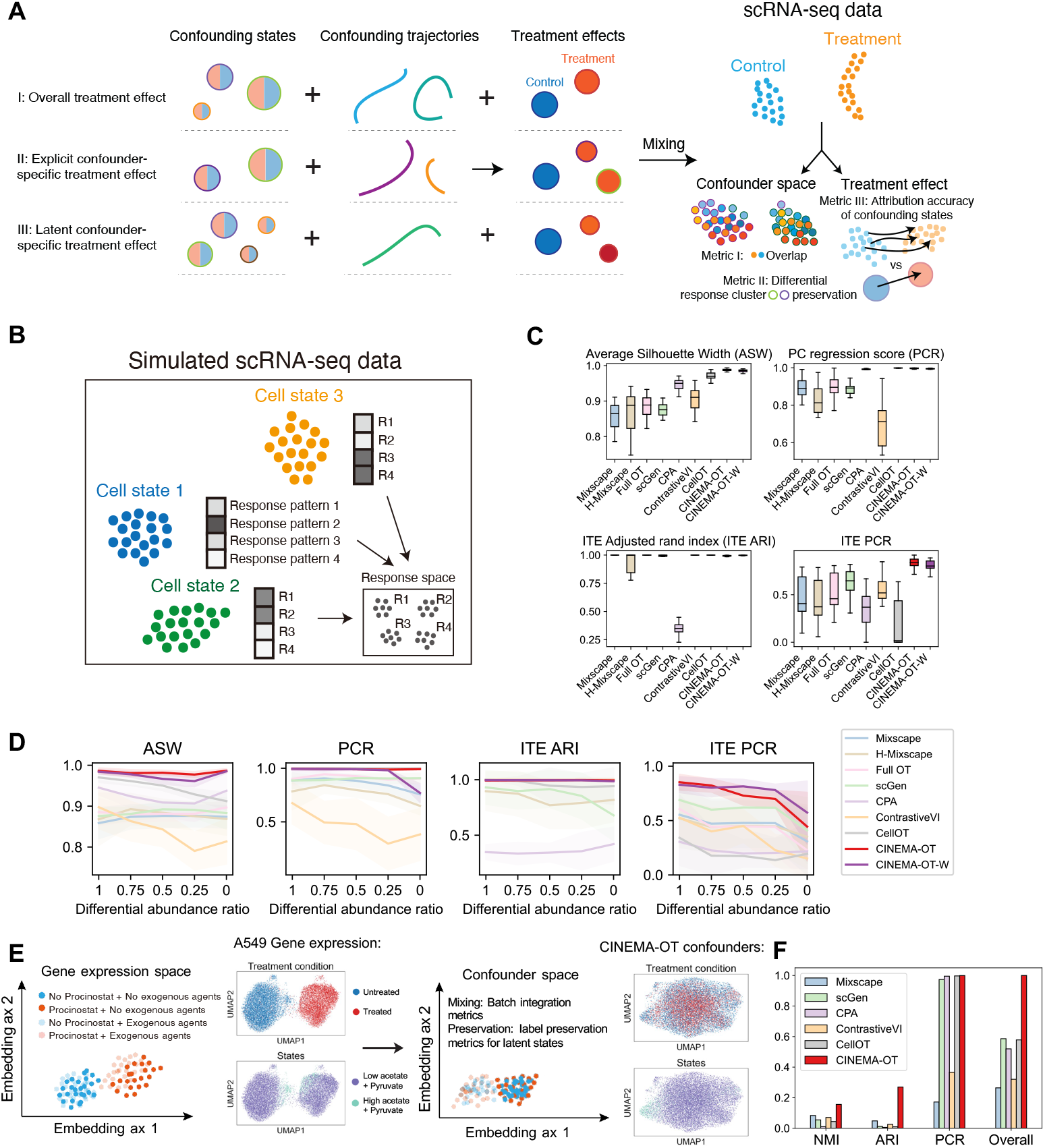
Benchmarking of CINEMA-OT against other methods for single-cell perturbation analysis. **A.** Illustrations of various factors to be considered in the task of single-cell level perturbation effect quantification. We evaluate CINEMA-OT against other methods using three classes of metrics: (I) mixing of samples in confounder space, (II) differential response cluster preservation in individual treatment response matrices, and (III) attribution accuracy of differential responses to confounding effects. **B.** Illustration of our conducted simulation study. **C.** Quantification of different validation metrics on synthetic data for CINEMA-OT and other methods. **D.** Comparison of different methods’ performance across synthetic datasets with various differential abundance (DA) ratio settings: 1, 0.75, 0.5, 0.25, 0 (Missing cell types). **E.** Illustration of the validation of CINEMA-OT on the Sciplex dataset. **F.** Quantification of different validation metrics on the Sciplex dataset for CINEMA-OT and other methods.

Prior to our evaluations, the optimal hyperparameter setting for each method was selected through parameter-sweep analysis (See Methods). Our quantitative assessment of these synthetic datasets shows that in the case of balanced confounder states (no differential abundance), CINEMA-OT(-W) achieves the best performances among all tested methods in batch mixing and treatment effect attribution, while most methods including CINEMA-OT(-W) succeed in differential response cluster preservation (Figure 3C).

By varying the differential abundance level, we find that the original version of CINEMA-OT performs better than CINEMA-OT-W when the differential abundance level is small (DA ratio ≥ 0.75), while CINEMA-OT-W performs better in treatment effect attribution at a higher differential abundance level (DA ratio ≤ 0.5). In both cases, CINEMA-OT significantly outperforms other methods in treatment effect attribution, while performing as well as other methods in cell distribution equalization and differential response cluster preservation (Figure 3D). The superior performance of CINEMA-OT-W in the datasets with significant differential abundance is also shown by qualitative visualizations of both confounder space (where the response cluster should be mixed and the cell states should be distinctive) and the treatment effect space (where the response cluster should be distinctive while the cell states should be mixed) (Supplementary Figure 6). Through our experiments with varying data sparsity levels (through subsampling), we have found that even under high sparsity level, CINEMA-OT’s performance only decreases slightly with increasing sparsity, maintaining a lead in accurately attributing responses while achieving top performances in preserving batch mixing and differential response clusters (Supplementary Figure 3).

Finally, we performed a benchmarking study of run time and peak memory usage on a series of subsampled scRNA-seq data, containing 1000, 2000, 5000, 10000, 20000, and 50000 cells. Our results show that CINEMA-OT and CINEMA-OT-W performs nearly as quickly as the fastest method available (Mixscape), significantly outperforming deep learning-based approaches in speed, with a runtime of approximately one minute for 50,000 cells (Supplementary Figure 4). Although CINEMA-OT(-W)’s peak memory usage is substantial due to the use of a dense matching matrix across conditions, it still requires less than 12GB of memory for 50,000 cells, making it feasible to run on most modern laptop computers (Supplementary Figure 4).

### 2.5 Validation of CINEMA-OT using real data

In order to evaluate the performance of CINEMA-OT in a real setting, we use two publicly available single-cell transcriptomics datasets: (1) sequencing of entorhinal cortex in patients with Alzheimer’s disease and unaffected controls [50]; (2) the sci-Plex4 drug perturbation dataset [8], which measures the response of the A549 and MCF7 cell lines to perturbation with 17 drugs.

In the Alzheimer’s disease dataset, we focus on qualitative comparison of perturbation effect removal and differential response cluster preservation. While the first comparison can be conducted in an unsupervised manner, for the second comparison we integrate prior knowledge to evaluate the preservation of clusters of interest [51]. One notable example gene selected is SPP1, which has been previously described as up-regulated in some cell types of patients with Alzheimer’s disease (e.g. microglia, some neuronal subtypes), but not in others (e.g. endothelial cells) [52, 50]. We compare CINEMA-OT to Mixscape, scGen, CPA, ContrastiveVI, and CellOT in our experiments, covering both the default model (cell-type unaware) and cell-type aware models for scGen and CPA. The visualizations of confounding spaces and treatment effects identified by each method can be seen in Supplementary Figure 8. Our results show that the other methods in general either preserves the differential response of SPP1 by automatic clustering (Mixscape, scGen, CPA without cell type label), or mix cell distributions well in the latent space (ContrastiveVI, CellOT), but not both. By contrast, CINEMA-OT succeeds in both tasks (Supplementary Figure 8-9).

In the Sciplex dataset, we investigate the response to perturbation with Pracinostat, a histone deacytelase (HDAC) inhibitor, with the combinatorial induction of exogenous acetate, citrate, and pyruvate. HDAC inhibitors act as antitumoral agents by antagonizing the pro-transcriptional effects of histone deacetylation and silencing the expression of oncogenic factors through chromatin remodeling [53]. As HDAC inhibitors act partly through the deprivation of acetyl-CoA, we expect that the relative abundance of acetyl-CoA precursors within a cell would modulate the effect of HDAC inhibitor exposure, and acetyl-CoA precursors can be considered confounders [8] (Figure 3E). Indeed, in the UMAP embedding of the A549 cell line gene expression across two Pracinostat doses, within each dose population, the cell neighborhood relationship is determined by doses of exogenous acetate, citrate, and pyruvate, separating the entire cell population into two latent confounder states. Ideally, a treatment-effect analysis method should not only achieve good mixing in the confounder space, but also automatically match the cells by the latent states in order to accurately specify the treatment effect. The two aspects can be quantitatively validated for each method by employing batch mixing metrics and label preservation metrics in the confounding space. Amongst all tested methods, CINEMA-OT achieves superior performance in both aspects, as suggested by our qualitative and quantitative evaluations (Figure 3F, Supplementary Figure 7).

### 2.6 CINEMA-OT identifies heterogeneous response patterns and synergistic effects of cigarette smoke exposure in rhinovirus infection

In addition to benchmarking CINEMA-OT against other methods, we have applied CINEMA-OT to new scRNA-seq data of rhinovirus infection in primary human bronchial organoids (Figure 4A). The experiment comprises 4 conditions: cigarette smoke extract (CSE) exposure, rhinovirus (RV) infection, the combination of rhinovirus and cigarette smoke (RVCSE), as well as a control condition (MOCK). While rhinovirus infection has been previously studied [54], the goal of our study was to probe cellular defense responses to viral infection from each airway epithelial cell type in the presence or absence of a common environmental insult known to impact the outcome of rhinovirus infection: cigarette smoke. Previous studies of viral infection using this model considered gene expression in each cell type but not heterogenous response patterns, which may be of biological and clinical relevance in understanding the tissue response to respiratory virus infections [54, 55].

**Figure 4:**
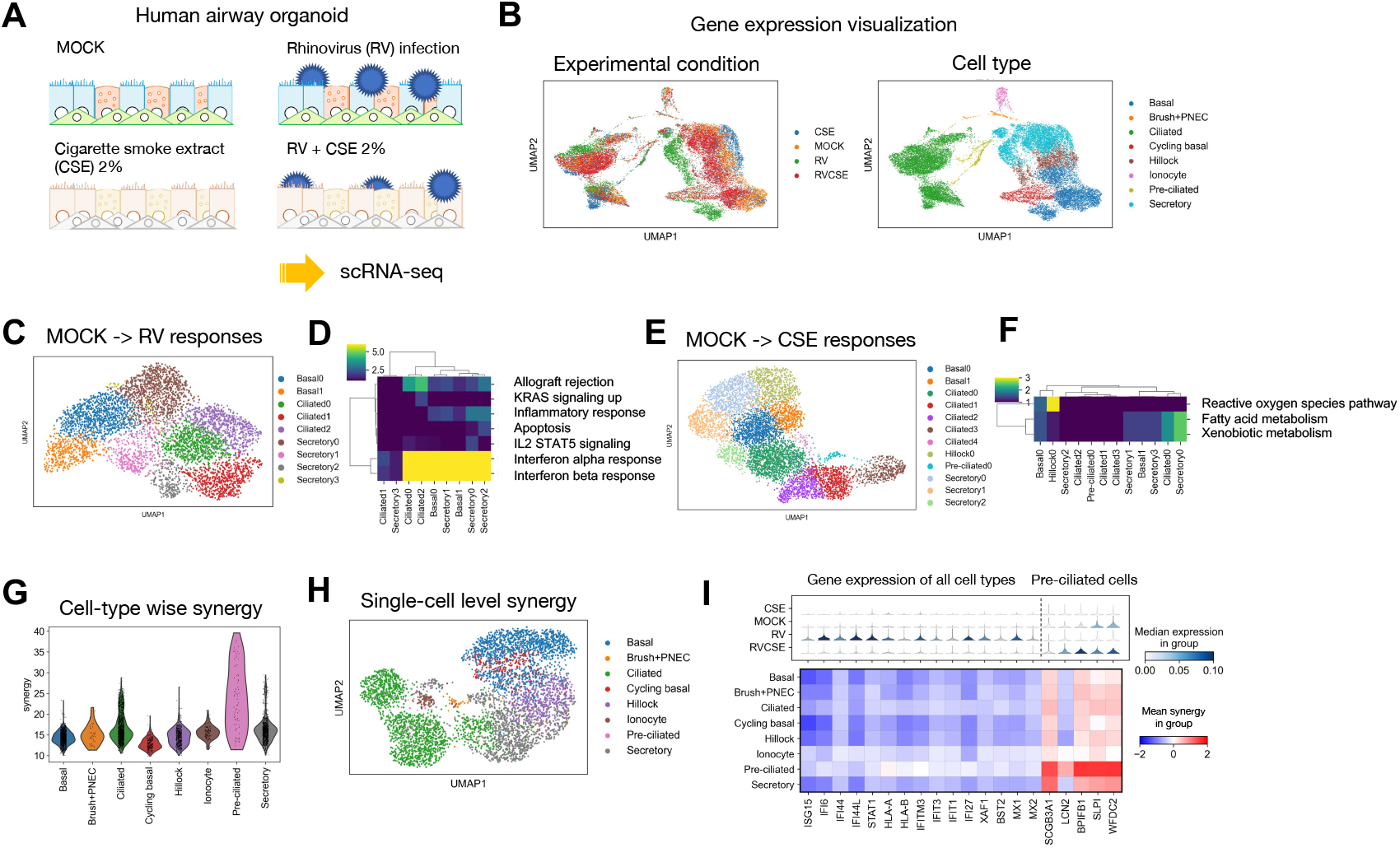
CINEMA-OT identifies a heterogeneous defensive response in human airway epithelial cells exposed to rhinovirus (RV) and cigarette smoke extract (CSE). **A.** Overview of experimental design. Differentiated airway epithelial organoids are challenged with mock (control) or RV 1A infection, with or without CSE exposure. **B.** UMAP projection of expression data labeled by perturbations and cell types. **C.** UMAP projection of the individual treatment effect matrix obtained by CINEMA-OT from the RV response without CSE exposure, colored by response clusters. **D.** Gene set enrichment analysis of MOCK to RV response clusters identified by CINEMA-OT. **E.** UMAP projection of the individual treatment effect matrix obtained by CINEMA-OT from the CSE response without RV exposure, colored by response clusters. **F.** Gene set enrichment analysis of MOCK to CSE response clusters identified by CINEMA-OT. **G.** Cell-wise synergy score visualization. **H.** UMAP visualization of CINEMA-OT synergy embedding, colored by cell types. **I.** Two diverging patterns among CINEMA-OT identified significantly synergistic genes, visualized by stacked violin plots and dotplots. The first pattern mainly consists of interferon stimulated genes globally attenuated in the RVCSE condition compared with the RV condition, leading to overall negative synergy. Genes in the second patterns are activated in pre-ciliated cells from MOCK and RVCSE conditions, leading to significant positive synergy.

We first performed preprocessing of the dataset and annotated 8 cell clusters in total, including major cell types in the airway (basal, secretory, ciliated) and other rare (ionocyte, PNEC, and brush) or transitional (hillock and pre-ciliated) cell types (Figure 4B). We then performed CINEMA-OT analysis on mock-RV and mock-CSE condition pairs to identify single-cell level treatment effects (the ITE matrix). As expected, in response to rhinovirus infection, most epithelial cell types exhibited robust induction of the interferon response with upregulation of several interferon stimulated genes, including ISG15, IFI44, STAT1, MX1, and others (Figure 4C, 4D). [54]. In response to cigarette smoke extract exposure, a subset of epithelial cells increased expression of genes associated with metabolism of reactive oxygen species (PRDX1, TXN) as well as fatty acid and xenobiotic metabolism (ADH1C, ALDH3A1, CYP1B1). Interestingly, responses to CSE were primarily enriched in particular cell subpopulations including hillock, ciliated, and secretory cells, in contrast to the global interferon response seen following rhinovirus infection (Figure 4E, 4F). This demonstrates a functional division of defense mechanisms in airway epithelium, with cell type specific responses to different insults.

After analysis of the effect of cigarette smoke and viral infection individually, positive and negative synergy between these two insults was assessed by calculating cell-gene synergy scores (Figure 4G, H). Among the most significant synergistic effects, we found that interferon stimulated genes (ISGs) exhibited negative synergy in general, showing a global reduction during viral infection in the presence of cigarette smoke compared to viral infection alone (Figure 4I), consistent with previous mechanistic studies from our group showing that the antioxidant defense response induced by CSE suppresses signaling pathways required for induction of interferon-stimulated genes in response to viral RNA in airway epithelial stem cells [56].

In addition to a global attenuation of the interferon response, we discovered that pre-ciliated cells, in particular, exhibit pronounced synergistic expression of a distinct set of genes when co-exposed to RV and CSE (Figure 4G, 4I). Pre-ciliated cells, sometimes referred to as “deuterosomal” cells, are developing multiciliated cells with the marker genes CCNO and CDC20B [57]. Pre-ciliated cells co-exposed to both viral infection and CSE show synergistic induction of genes encoding secreted proteins that are typically associated with secretory cells in resting cultures, including SCGB3A1, LCN2, BPIFB1, SLPI, and WFDC2 (Figure 4I). This pattern could arise from pre-ciliated cells adopting a more secretory phenotype during co-exposure, or secretory cells adopting a pre-ciliated phenotype. These findings highlight the use of CINEMA-OT to identify synergistic effects on gene expression induced by co-exposure to viral infection and cigarette smoke.

### 2.7 CINEMA-OT reveals principles of innate immune response modulation from combinatorial interferon stimulation

Type I, type II, and type III interferons (IFNs) act as central regulators of immune responses during intracellular pathogen infection, cancer, and in auto-immunity. However, despite the identification and adoption within the literature of a core set of interferon-stimulated genes (ISGs), IFN responses can vary widely by cell type, by individual, by IFN stimulus type, by chronicity of exposure, and by combination with signals delivered by other cytokines. In other words, the interferon response is highly context dependent. This complexity, heterogeneity, and context-specificity of IFN signaling can lead to counter-intuitive results. For example, IFN*γ* is proposed to play both stimulatory and suppressive roles in cancer, and type I IFNs are used both as an immunosuppressant to treat multiple sclerosis and as immunostimulatory adjuvant treatments for cancer (e.g. melanoma) and chronic viral infection (e.g. HCV) [58–60]. To characterize the complexity of IFN signaling, we conducted acute (2 days) and chronic (7 days) stimulations of peripheral blood mononuclear cells (PBMCs) from multiple healthy donors with type I, type II, and type III IFNs, separately as well as in combination with other cytokines such as TNF*α* and IL-6 (Figure 5A-B).

**Figure 5:**
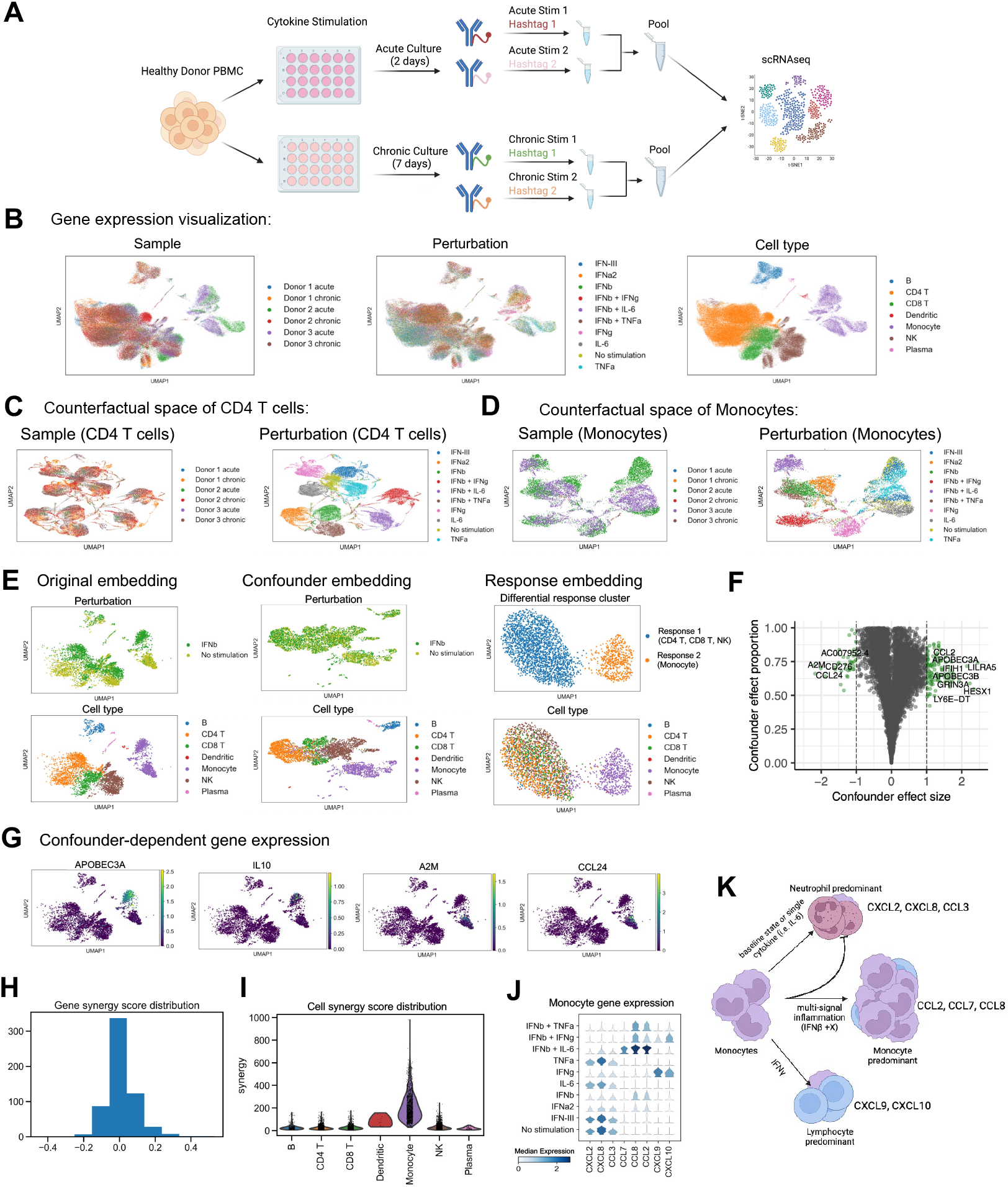
CINEMA-OT reveals combinatorial mechanisms of acute and chronic cytokine stimulation. **A.** Illustration of experimental design. **B.** UMAP projection of expression data colored by samples, perturbations, and cell types. In sample labels, H refers to the donor number, and D refers to the number of days of stimulation. **C.** UMAP projection of the CD4^+^ T cell counterfactual space from CINEMA-OT. Projections are colored by experimental batch and perturbation type. **D.** UMAP projection of the monocyte counterfactual space from CINEMA-OT. Projections are colored by experimental batch and perturbation type. **E.** UMAP projections of original data, confounder embedding, and individual treatment effects identified by CINEMA-OT respectively after acute stimulation with IFN*β* in H3D2, colored by response cluster and cell type. **F.** Volcano plot highlighting genes with significant confounder-specific treatment effect. **G.** Representative confounder-specific treatment-associated gene expression in original UMAP space. **H.** Gene synergy score distribution combining IFN*β* + TNFα, IFN*β* + IFN*γ*, and IFN*β* + IL-6 combinatorial treatment for in H3D2. **I.** Cell-wise synergy score visualization in acute condition, taking a single experimental batch (H3D2) as an example. **J.** Stacked violin plot of significantly synergistic chemokines identified by CINEMA-OT. **K.** Patterns of chemokine secretion programmed by single or multi-signal cytokine stimulation.

In order to understand the underlying structure of PBMC cellular response to interferon stimulation, we use CINEMA-OT to match treatment conditions to the untreated (control) condition. This analysis highlights the underlying hierarchical structure of cellular responses. As the hierarchical structure of cytokine response can vary with cell types, we pool CD4^+^ T cells and monocytes across experimental batches and run CINEMA-OT analysis. In this case, a confounder is defined by each different experimental batch. In CD4^+^ T cells, the response can be characterized by four meta-perturbation clusters: No stimulation, IFN*γ*, IL-6, TNF*α*; IFN*α*2, IFN*β*, IFN*β*+TNF*α*; IFN*β*+II.-6; IFN*β*+IFN*γ* (Figure 5C). In monocytes, a similar structure is observed, except that IFN*γ* in monocytes represents a distinct response cluster in the phenotypic space (Figure 5D).

Next, to demonstrate CINEMA-OT’s power in general single-cell level treatment analysis, we focus on analyzing the treatment effects of IFN*β* in a single experimental batch (H3D2). CINEMA-OT analysis highlights the induction of coordinated immune responses across cell types as well as celltype specific responses using a confounder-specific effect volcano plot. For example, despite a global change in interferon stimulated genes (Supplementary Figure 14), monocytes demonstrate a unique program characterized by increased APOBEC3A and IL10 and decreased A2M and CCL24 compared to other cell types (Figure 5E-G, Supplementary Figure 14). Notably, a similar qualitative analysis to which we performed on the Alzheimers dataset shows that CINEMA-OT achieves both good batch mixing and reasonable response clustering, compared with alternative tested methods (Supplementary Figure 12). CINEMA-OT is also applied to understand the treatment effects in CD4^+^ T cells of chronic versus acute stimulation and reveals the attenuation of genes involved in the core type I IFN response (Supplementary Figure 15).

To estimate the synergistic effects of acute combinatorial cytokine stimulation, we used CINEMA-OT to calculate cell-gene synergy scores. We next performed gene synergy score analysis by computing the gene-wise synergy score (See Methods). The gene synergy score analysis identified genes that were synergistically induced by each combinatorial perturbation (Figure 5H). Based on selected significant synergy genes, we summarized the cell-wise synergy effect by taking the norm over selected synergy genes. We have found that monocytes exhibit the most significant synergistic regulation compared with other cell types (Figure 5I). Further enrichment analysis identifies a number of chemokines with specific synergistic expression in monocytes with respect to different interferon perturbation (Figure 5J, Supplementary Figure 16). These chemokines exhibit synergistic patterns of response to multi-signal inflammation (e.g. IFN*β* + IL-6) in monocytes including inhibition of baseline neutrophil chemotactic signaling and induction of a monocyte chemotactic signaling program. The addition of IFN*γ* contributes lymphocyte predominant chemokines, while maintaining this core inflammatory programming (Figure 5J, 5K). These results suggest that CINEMA-OT, when applied to combinatorial experiments, is capable of revealing the synergistic logic governing cellular signaling in inflammation and tissue repair.

## 3 Discussion

With rapidly developing high-throughput screening technologies and an ever rising number of datasets, single-cell level experimental effect analysis is becoming a critically important task in biological discovery. Current analytical approaches aiming to address this need face a number of challenges. When treatment effects are confounder specific and do not change relative cell proportions, differential abundance methods may be unsuitable for extracting the dependence between confounder states and treatment responses. Recent neural network-based methods for characterizing perturbation effects learn nonlinear interactions between confounders and treatment effects, but may be prone to overfitting and have limited interpretability. Moreover, none of the existing approaches employ a causal inference framework for perturbation analysis.

In response to these challenges, we present CINEMA-OT, a framework for single-cell causal treatment effect analysis. By explicitly separating confounder and treatment signals and matching at the single-cell level, CINEMA-OT produces a per-cell view into the effects of experimental perturbations and conditions including disease states.

We applied CINEMA-OT in several use cases including synthetic and real data sets. In benchmarks, CINEMA-OT was able to outperform other methods in experimental perturbation analysis. In human airway organoids, CINEMA-OT revealed how cigarette smoke exposure can interfere with the normal innate immune response to rhinovirus infection. In combinatorial cytokine stimulation of *ex vivo* human peripheral immune cells, CINEMA-OT revealed complex logic which may underlie the specific recruitment of cells from the periphery to tissues responding to various injuries.

Two potential challenges for CINEMA-OT can arise due to bias-variance tradeoffs in optimal transport and the magnitude of batch effect versus biological perturbation effect. For the first challenge, a large smoothness threshold in the entropy regularized OT method can overly smooth the obtained matching and cause false positives by incorrectly identifying confounder variation as treatment-associated variation. However, too small a threshold would both harm the method’s stability and cause high variance. In practice, an adequate threshold can be chosen based on coarse-graining of the matching matrix. For the second challenge, in the CINEMA-OT framework, as CINEMA-OT performs matching in the confounder space, the CINEMA-OT identified confounding space and the optimal transport matching plan are minimally altered by the level of batch effect, as the batch effect can be viewed as a treatment-induced factor itself. However, at the gene level, as the current implementation of CINEMA-OT does not perform count modeling, the differential expression analysis may be still affected by the batch effect when it causes significant distortions of global gene expression. In this case, the confounder matching identified by CINEMA-OT still provides meaningful neighborhood relationships for conducting differential expression testing [61]

CINEMA-OT is designed to estimate causal responses from experimentally perturbed single cell omics measurements. CINEMA-OT is not able to extrapolate, meaning it cannot identify the causal effect of unmeasured perturbation-cell pairs. Integrating prior knowledge (i.e: ChemCPA [62], expiMap [63]) to achieve causally-meaningful extrapolation for unseen perturbation effects remains a promising future direction to explore. Moreover, while we have implemented an reweighting procedure to account for differential confounder abundance that may arise in response to treatment, CINEMA-OT is not designed for cases where changes to confounder distributions are the primary effects of interest. In those cases, tools such as MELD, MILO, or DA-seq may be more suitable [17, 14, 15].

As a highly explainable and scalable causal framework, we anticipate CINEMA-OT to be widely adopted for single-cell perturbation analysis.

## Data availability

The data of Sciplex is taken from the original publication [8] and the processed Alzheimer data is accessed from ContrastiveVI’s tutorial: https://drive.google.com/uc?id=1R1aN-LWUQ6c_N44n5-xjy2nEPzl7H0Dc. We provide the public access to a subset of combinatorial cytokine simulation data: https://drive.google.com/file/d/1A3rNdgfiXFWhCUOoUfJ-AiY7AAOU0Ie3/ view. The other newly produced datasets (rhinovirus infection scRNA-seq data, the full dataset of interferon stimulation scRNA-seq profiling) are available upon reasonable request.

## Code availability

CINEMA-OT is implemented as a public available open-source Python package, available at https://github.com/vandijklab/CINEMA-OT.

## Methods

### CINEMA-OT

CINEMA-OT is an unsupervised method for separating confounding signals from perturbation signals for matching cells via imputing counterfactuals and computing perturbation effect at a singlecell level (https://github.com/vandijklab/CINEMA-OT). The detailed workflow of CINEMA-OT is as follows.

1. **Rank initialization** In order to perform CINEMA-OT, we first need to initialize the expected matrix rank, representing the total signal number. We here offer two possible approaches for rank initialization in CINEMA-OT. Biwhitening [64] is a recently-developed method to remove independent heteroskedastic noise in data with inspirations from random matrix theory. It does diagonal matrix transformation of the data on both sides and thresholding based on the Marchenko-Pastur law [65]. After thresholding, we can get the true matrix rank and the matrix’s low dimensional approximation. Mathematical details of biwhitening can be seen in [64]. In CINEMA-OT, we have implemented a version of biwhitening with fixed hyperparameters. In large datasets, we suggest using prespecified rank values. Empirically, we have found that CINEMA-OT is robust to rank selection at certain ranges and can give a good performance when DimSize ∈ [20,50].
2. **Signal selection with independent component analysis** Independent component analysis is already a well-addressed method in data analysis and has various implementations. Here we use the FastICA implementation from the package `sklearn.decomposition` [66]. Prior to FastICA, input data is PCA-transformed using scanpy [67]. In order to identify confounder signals and treatment-associated signals, we have adopted a recently proposed cross rank coefficient [30], which is able to quantify the functional dependence between ICA signals and query signals (in this case, the treatment signals). We use the implementation of this method from the XICOR package in Python. The threshold of the cross rank coefficient is set to be 0.05 to 0.75 in this study. We note that tuning the threshold parameter has a practical meaning in the algorithm. High thresholds correspond to less tolerance for false positive treatment signals, which leads to local matching more similar to Mixscape analyses. Meanwhile, setting a low threshold means less tolerance for false positive confounder signals and can lead to lower resolution of matching, which, in the extreme case, coincides with pseudo-bulk differential expression testing methods.
3. **Optimal transport matching** After selecting confounding signals, we perform matching across treatments via optimal transport, which provides a smooth transport map and does not require neighbor number selection. Here we consider the entropy regularized optimal transport formulation, which can be efficiently solved by the Sinkhorn-Knopp algorithm [31]. In this formulation of the problem, the penalty coefficient act as a hyper parameter influencing the resolution and smoothness of the transport map. We have empirically determined that the optimal value for the penalty coefficient often lies within the range (10^-6^, 10^-3^) multiplied by the number of confounding signals.

#### Algorithm 1 CINEMA-OT

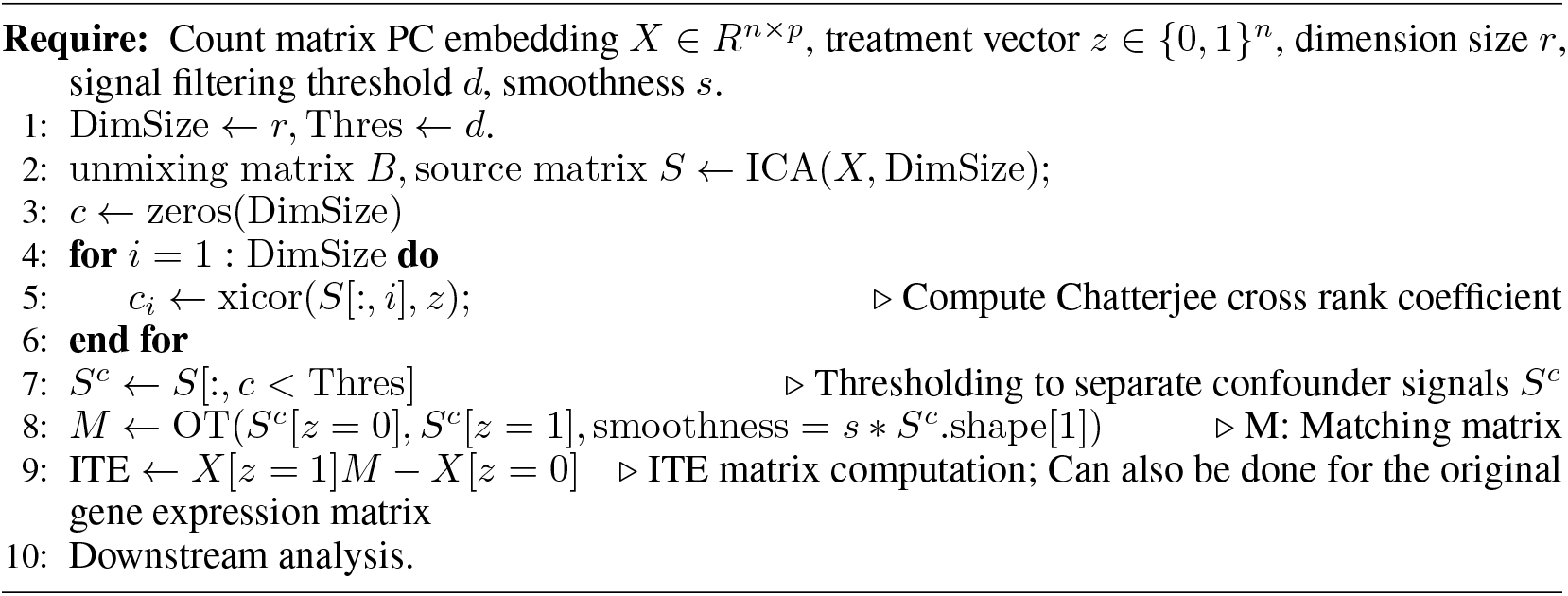

#### Algorithm 2 OT

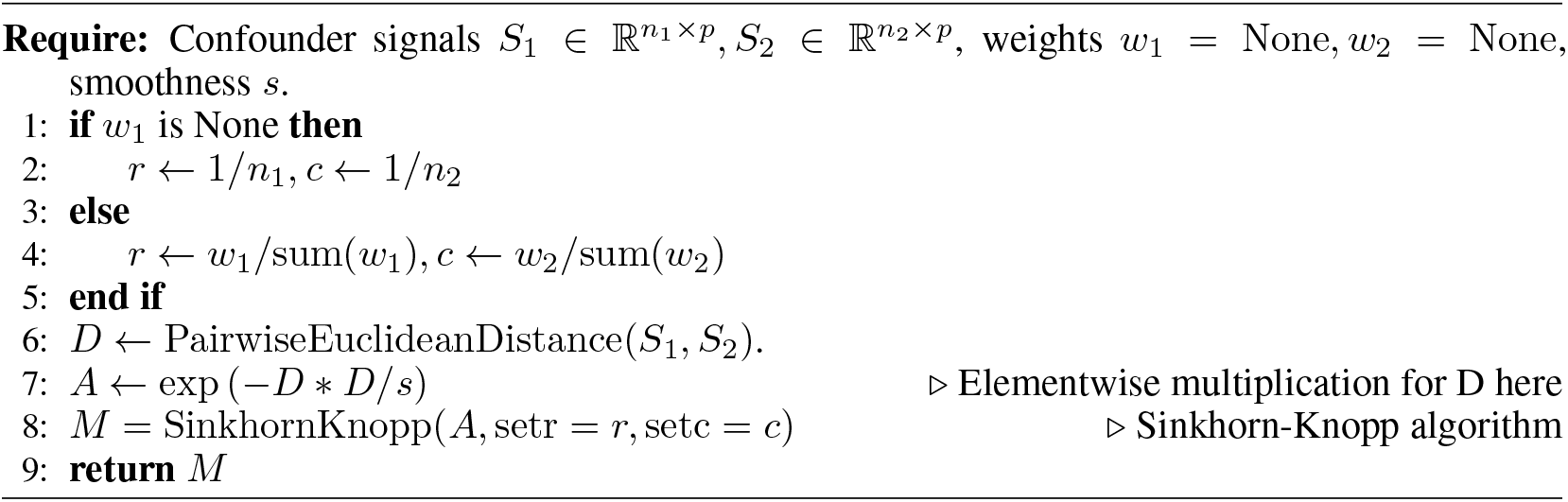

### CINEMA-OT-W

In CINEMA-OT-W, the treated cells are first aligned by their 20 nearest neighbors in the untreated condition. Then Leiden clustering is performed on the full aligned cell set at a pre-specified resolution (`r`). Foe each Leiden cluster, the cells from one of the conditions are subsampled such that the number of cells are the same for each condition in the same cluster. After subsampling, the confounder signals are independent of treatment event. Therefore, the ICA procedure can be conducted on the subsampled data, and the confounder component selection is performed on the identified independent components. The entropic regularized OT can be later performed on the confounder components of either subsampled data or the full data. In the latter case, the ICA unmixing matrix computed from subsampled data is applied on the full data embedding. Notably, as control state indicates the cells in the normal states in most cases, we may assume that the untreated cells always covers the confounding states of the treated cells. In this case, the treatment effect of all treated cells can be computed by CINEMA-OT-W.

In pratical datasets, the underlying confounders are often provided in terms of cell-wise labels (such as cell types), which indicates biological meaningful sampling labels. Therefore apart from CINEMA-OT-W, CINEMA-OT offers users the ability to specify known confounder labels (e.g. cell type, cell cycle), without the need for a sampling procedure.

### Downstream analysis

1. **Visualization and clustering of the ITE matrix** With the ITE matrix computed by matching counterfactuals, we are able to numerous standard analyses. We may employ dimensionality reduction techniques such as t-SNE, UMAP, or PHATE [68–70] to visualize clusters in the response space. We may also employ clustering techniques, such as Leiden clustering [23] to group cells by similarity of treatment responses.
2. **Synergy analysis** For synergy effect, we compare ITE matrices for two treatment conditions against the ITE matrix for the combined treatment. Formal derivation of the synergy score is given as follows. Consider *D*_*A*=í1,*B*=0_ as the ITE matrix for treatment *A* alone, *D*_*A*=0,*B*_=1 as the ITE matrix for treatment *B* alone, and *D*_*A*=1,*B*=1_ as the ITE matrix for the combined treatment. We may define a synergy matrix Ψ as:

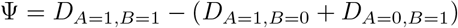 Where each entry Ψ*_g,c_* represents the synergy score for gene *g* and cell *c*. In order to test if a particular gene *g* has significant synergistic effect, we formulate the problem as if we should reject

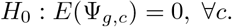 Note here if we apply no normalization, we are aiming for additive synergy; if we instead apply log normalized data, *H*_0_ would test for multiplicative synergy. We assume that different cells are unlikely to have opposite synergy effects, allowing us to relax *H*_0_ as:

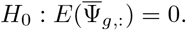 Assume the new *H*_0_ holds, then for each gene *g*, we compute the empirical synergy:

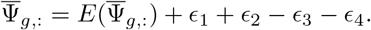 Because here *H*_0_ holds, the expectation of the noise term is zero. Assume the noise is Poisson, then with the property of Poisson distribution, in the case of log normalization, *ϵ_i_s* are averages of i.i.d. scaled log1p Poisson distribution with zero expectation. With the delta method, the variance of the noise term is approximated as:

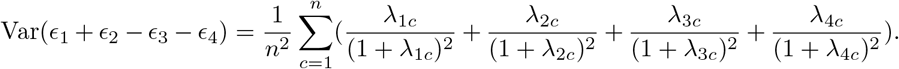 Where λ*_ic_*, *i* ∈ {1, 2, 3,4}s are counterfactual cell gene expression expectation for each cell in 4 conditions. We note the formula 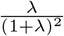 is self-standardized as it is a smooth function and is near zero for either large or small λs. Therefore, to simplify the statistical test, we assume

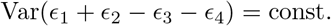 In this case, if we define

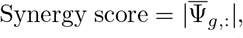

identifying most synergistic genes among all genes can be turned into comparing the synergy score over all genes.
3. **GSEA analysis** For differential gene expression significance, we have applied the non-parametric Wilcoxon signed-rank test. We apply a p-value threshold (10^-5^) and expression fold change threshold selected by the user to identify significantly regulated genes. These genes are input into GSEApy for analysis by functional signatures [71, 72].
4. **Attribution analysis** By clustering cells both by treatment responses (i.e. using the ITE matrix) and control condition clusters (i.e. cell subtypes), the matching matrix from CINEMA-OT can be coarse grained. The resulting course-grained matching matrix is of shape `ResponseClusterNumber × ControlClusterNumber`. Each column of the matrix gives the likelihood of a control condition cluster to have different modes of response. By reading each row of the matrix, we are able to attribute each response to the underlying control condition cluster. Furthermore, to investigate the genes with significant confounder-specific treatment effect, we fit each gene to the causal regression model

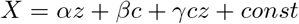 In this case, the confounder-specific effect part is *γcz*, whose significance can be established by classical linear regression theory. However, in our case, the noise term can stand for latent-factor specific effect thus not satisfying the assumption of classical regression. Therefore, here we instead quantify the ratio of norm between confounder-specific effect and the residual as an indicator of confounder-specific effect significance.

#### Algorithm 3 CINEMA-OT-W

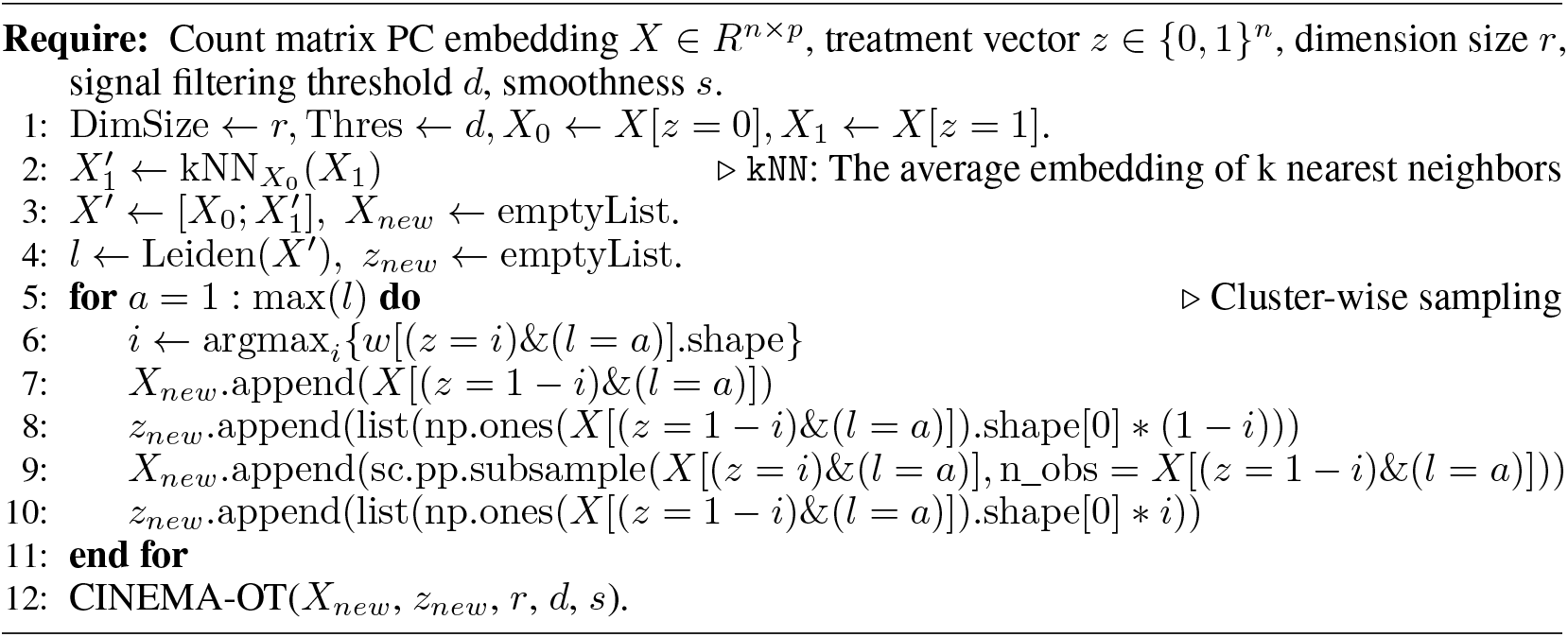

### Data simulation and analysis

For the simulation in main figure (Figure 2B-C), we used Scsim to simulate 1000 gene by 5000 cell count matrices with 2-5 underlying cell states with 2 gene regulation programs. For each cell state, we simulated a random discrete distribution to represent the corresponding response distribution of the cell state. Then the response count matrix of shape 500 gene times 5000 cells were simulated and concatenated with the confounder count matrix.

For the Mixscape analysis, we have implemented a simple version in Python that matches cells across conditions according to the descriptions in [18]. For Harmony-Mixscape analysis, we have used the Python package `harmonypy` (https://github.com/slowkow/harmonypy) with default settings [73], and apply Mixscape on the batch corrected embeddings returned by Harmony. For Full OT analysis, we implemented a function that calls entropy-regularized optimal transport on the full PC embedding space with a tunable smoothness parameter. For scGen, CPA, ContrastiveVI and CellOT, the default model settings were used consistent with those provided in their tutorials respectively (https://scgen.readthedocs.io/en/stable/tutorials/scgen_perturbation_prediction.html (scGen), https://github.com/facebookresearch/CPA/blob/main/notebooks/demo.ipynb (CPA), https://colab.research.google.com/drive/1_R1YWQQUJzgQ6kz1XqglL5xZn8b8h1TX?usp=sharing (ContrastiveVI), https://github.com/bunnech/cellot/blob/main/configs/models/cellot.yaml (CellOT)). For CellOT, we inputted principal component embeddings for training and evaluation.

In order to investigate the effect of hyperparameter settings on different methods respectively, we performed parameter-sweep analysis for all tested methods. The sweeped hyperparameters for all methods are summarized as follows:

- Mixscape (The number of neighbors `k`): [5,10,20 (Default),50,100];
- Harmony-Mixscape (The number of neighbors `k`): [5,10,20 (Default),50,100];
- Full OT (Epsilon e): [0.1,0.3,1,3];
- scGen (KL weight *l*): [0,5e-6,5e-5 (Default), 5e-4];
- CPA (Adversary strength *l*): [5,20,60 (Default), 200];
- ContrastiveVI (Wasserstein penalty *l_MMD_*): [0 (Default),1e-4,1e-3,1e-2,1e-1];
- CellOT (Frobenius norm regularization reg): [0.01,0.1,1 (Default),10];
- o CINEMA-OT (confounder threshold `cutoff`): [0.05,0.1,0.15,0.2,0.25]; (OT smoothness e based on the optimal `cutoff`): [1e-5,3e-5,1e-4,3e-4,1e-3];
- CINEMA-OT-W (Leiden clustering resolution r based on optimal parameters of CINEMA-OT): [0.3,0.6,1,1.21.

The parameter-sweep analysis results are listed in Supplementary Figure 2. Based on the four metrics, the hyperparameter settings used throughout our benchmarking analysis were selected: `k` = 20 (Mixscape); `k` = 20 (Harmony-Mixscape); `e` = 0.1 (Full OT); *l* = 0 (scGen); *l* = 20 (CPA); *l_MMD_* = 0 (ContrastiveVI); `cutoff` = 0.05, `e` = 1e-5 (CINEMA-OT); `r` = 0.6 (CINEMA-OT-W).

Based on the optimized hyperparameters, a number of following analyses were performed:

1. **The effect of differential abundance on single-cell treatment effect analysis.** To explore the effect of differential abundance on the performance of single-cell treatment effect analysis methods, we selectively subsampled cells from half of the confounder clusters in the treated condition. The subsample ratio, which we refer to as differential abundance ratio, are selected as different levels: [1,0.75,0.5,0.25,0]. The case of DA ratio = 1 corresponds to no differential abundance effect and the zero case corresponds to the case that certain cell populations are not observed in the treated condition.
2. **The effect of noise level on single-cell treatment effect analysis.** To perform the analysis, the count matrix were subsampled according to the Scanpy function `sc.pp.downsample_counts` with the `total_counts` parameter specified to be (1,0.5,0.2,0.1,0.05) fold of the original count matrix’s total count number.
3. **Running time and peak memory usage.** We conducted the scalability analysis by testing different methods’ run time and maximum memory usage on subsampled interferon datasets, with cell numbers ranging in [1000,2000,5000,10000,20000,50000]. For Mixscape, scGen, CPA, and CINEMA-OT, the data was normalized and log transformed, and we selected 773 highly variable genes using mean ? dispersion thresholds provided by the default Scanpy function `sc.pp.highly_variable_genes(adata, min_mean=0.0125, max_mean=3, min_disp=0.5)`. In the case of Contrastive-VI, which models the distribution of the count matrix, we used the original count matrix of highly variable genes.

### Benchmarking metrics

Average Silhouette width (ASW), Principal component regression score (PCR), and adjusted Rand index (ARI) are batch mixing and biological preservation metrics used to evaluate batch correction methods performance in the systematic benchmarking paper [51]. CINEMA-OT uses the first two metrics to evaluate mixing in confounder space, as a surrogate for correct matching that can still be measured when ground truth labels are not present. ARI is used to evaluate the preservation of response clusters in ITE matrices estimated from simulated data. The PCR for ITE matrices is used to evaluate attribution accuracy of response, as in our experimental settings the response of each cell is conditionally independent of cell states conditioning on the response cluster assignments. For all metrics, we use the implementations of these metrics from package scib [51].

### Alzheimers scRNA-seq data

The Alzheimer scRNA-seq data was downloaded from https://drive.google.com/uc?id=1R1aN-LWUQ6c_N44n5-xjy2nEPzl7H0Dc. For Mixscape, scGen, CPA and CINEMA-OT, the data was normalized and log transformed. 2000 highly variable genes were selected with Seurat v3 approach implemented in Scanpy. As Contrastive-VI models the distribution of count matrix, the original count matrix of highly variable genes was used for Contrastive-VI. For CellOT, we inputted principal component embeddings computed from preprocessed highly variable genes for training and evaluation.

### Sci-Plex4 data

The Sci-Plex4 data was accessed from https://www.ncbi.nlm.nih.gov/geo/query/acc.cgi?acc=GSM4150379 with GEO accession number GSM4150379. The data is preprocessed via protocol https://github.com/manuyavuz/single-cell-analysis/blob/main/single_cell_analysis/datasets/sciplex.py. After preprocessing, we normalized and log transformed the raw count matrix, then performed highly variable gene selection using mean / dispersion thresholds provided by the default Scanpy function `sc.pp.highly_variable_genes(adata, min_mean=0.0125, max_mean=3, min_disp=0.5)`. Finally, we performed subsequent analysis described in main text for Mixscape, scGen, CPA and CINEMA-OT. The original count matrix of highly variable genes was used to evaluate ContrastiveVI. For CellOT, we inputted principal component embeddings computed from preprocessed highly variable genes for training and evaluation.

After estimating all metrics, each metric was rescaled so that the max value all methods tested equals 1. Then we computed the average of batch mixing score (PCR) and label preservation score (the average of NMI and ARI) as the final metric used (Overall_score).

### Rhinovirus infection data

Primary human bronchial epithelial cells from healthy adult donors were obtained from commercial vendor (Lonza) and cultured at air-liquid interface according to the manufacturers instructions (Stem Cell Technologies) using reduced hydrocortisone. Cells were kept at air-liquid interface for 4 weeks before experiment; maturation of beating cilia and mucus production was confirmed using light microscope. Cells were then infected with mock or 10^5^ PFU human rhinovirus 1A per organoid, with or without exposure to 2% cigarette smoke extract (CSE). Single cell suspension is collected by trypsin digestion at 5 days post infection and submitted to single cell RNA sequencing using The 10X Genomics single-cell 3’ protocol. The final dataset contains 26420 cells in 4 samples (mock, RV, CSE, RVCSE). We performed normalization (by `sc.pp.normalize_total`), log1p transformation, highly variable gene selection using mean ? dispersion thresholds provided by the default Scanpy function `sc.pp.highly_variable_genes(adata, min_mean=0.0125, max_mean=3, min_disp=0.5)`, scaled their values for PCA and Leiden clustering analysis. We annotated 8 cell clusters based on known cell type markers of airway epithelial cells [74]: cycling basal, basal, hillock, secretory, pre-ciliated, ciliated, ionocyte, and PNEC/brush cells. CINEMA-OT analysis on mock-RV and mock-CSE was run with default parameters with `smoothness=1e-5`. Synergy analysis was performed with `smoothness=3e-5`.

### Interferon treatment data

#### PBMC processing and in vitro culture

The study was approved by Institutional Review Boards at Yale University (following Yale melanoma skin SPORE IRB protocol). Healthy donors consented to donation of peripheral blood for research use.

Human PBMC were isolated using Lymphoprep density gradient medium (STEMCELL). PBMC were plated at 1 million cells per ml and stimulated with 1000U/ml human IFN*α*2 (R&D systems), 1000U/ml human IFNβ (pbl assay science 11415), 1000U/ml human IFN*γ* (pbl assay science), 1ug/ml human IFN-III/IL-29 (R&D systems), 100ng/ml human IL-6 (NCI Biological Resources Branch Preclinical Biologics Repository), 20ng/ml human TNFα (R&D systems), and combinatorial cytokines IFN*β* + IL-6, IFN*β* + TNFα, IFN*β*+ IFN*γ* at indicated concentrations above for up to 48 hours.

#### Cell enrichment and 10x sample preparation

Cultured cells were collected stained with TotalSeq anti-human hashtags C0251-C0260 (Biolegend), viability dye (zombie red, Biolegend) and anti-human CD45-FITC (clone HI30, Biolegend) and enriched for live CD45^+^ cells using BD FACS Aria II. Sorted cells were then resuspended to 1200 cells per ul and barcoded for multiplexed single cell sequencing using 10x Genomics 5v2 chemistry (10x Genomics, PN-1000263).

#### Sequencing and 10x sample alignment

Single cell RNA sequencing libraries were sequenced on Illumina NovaSeq at read length of 150bp pair-end and depth of 300 million reads per sample.

#### scRNA-seq data analysis

Data from three donors across Day 2 and Day 7 were concatenated together into labeled anndata objects for analysis. For each of the 6 samples, we filtered cells with less than 200 genes and we filtered genes expressed in fewer than 3 cells. For further quality control, cells with a high proportion of mitochondiral reads (> 7%) were excluded. The distribution of genes per cell was visually inspected and upper thresholds selected on a persample basis to exclude doublets. For each of the samples, the upper threshold was selected as [6000,3500,4000,3500,4500,3500] respectively. Following filtering, the count data was normalized and log transformed. Highly variable gene were selected using mean / dispersion thresholds provided by the default Scanpy function `sc.pp.highly_variable_genes(adata, min_mean=0.0125, max_mean=3, min_disp=0.5)`. Highly variable genes were scaled for subsequent PCA and UMAP projection.

For individual treatment effect analysis, we additionally filter T cell receptor genes, histocompatibility genes, and immunoglobulin genes from the highly variable gene set. Genes to be filtered were obtained from the HUGO database [75]. After filtering, highly variable genes were used for downstream visualization analysis.

CINEMA-OT analysis was run on each of the samples separately, with `thres=0.5, smoothness=1e-4, eps=1e-2`, and preweights given by cell types. The implementation of other methods were consistent with the experiments conducted in the Sciplex dataset.

For the synergy analysis of donor 3 on day 2 (H3D2), we selected significant synergy genes by a absolute value threshold of 0.15.

