## Supplementary Material for "Causal identification of single-cell experimental perturbation effects with CINEMA-OT"

Preprint version.

### 1 Supplementary notes

**Theoretical foundation of the CINEMA-OT framework**

Here we give an rigorous treatment of the causal framework and underlying assumptions in CINEMA-OT.

> **Assumption 1 (Formal): Independent sources and noise.** *Confounding factors and treatment events $(s_1, ..., s_l, z)$ are independent random variables. The treatment event $z \in \{-1, 1\}$ with $P(z = 1) = p$.*
>
> **Assumption 2 (Formal): Linearity of source signal combinations.** *The expression of each gene can be linearly decomposed as the mixing of noisy confounding signals and treatment-associated signals. Without loss of generality, we assume the concatenated random vector $(s_1 + e_1, ..., s_l + e_l, z + e_z)$ is whitened and at most only one of the $(s_1 + e_1, ..., s_l + e_l, z + e_z)$ is Gaussian. The observed data matrix $X \in \mathbb{R}^{n\times d}$ is generated by mixing of i.i.d sampled noisy signals plus i.i.d noise terms $\epsilon$.*

In our formulation, to be more realistic, we consider both the biological variation terms $e$ and the measurement noise $\epsilon$. For simplicity, we may understand the signal $s$ as cell types and $e$ as biological variations contributing to the heterogeneity within each cell type. Given assumption 1 and 2, denote the gene count matrix as $X \in \mathbb{R}^{n\times d}$, and it is generated by a random vector $\boldsymbol{x} \in \mathbb{R}^m$ with the mixing matrix $A$. Then the data generation mechanism is given by

$$\boldsymbol{x}^i \overset{\text{i.i.d}}{\sim} \begin{bmatrix} s_1 + e_1 \\ \vdots \\ s_l + e_l \\ z + e_z \end{bmatrix}; \quad \boldsymbol{\epsilon}^i \text{ i.i.d;} \quad X = [\boldsymbol{x}^1, \boldsymbol{x}^2, ..., \boldsymbol{x}^n]^T A + [\boldsymbol{\epsilon}^1, \boldsymbol{\epsilon}^2, ..., \boldsymbol{\epsilon}^n]^T. \tag{1}$$

As $n \to \infty$, by the subspace consistency established in [1], we have the first $l+1$ identified principal components of form:

$$\hat{\boldsymbol{x}} \overset{\text{i.i.d}}{\sim} B \begin{bmatrix} s_1 + e_1 \\ \vdots \\ s_l + e_l \\ z + e_z \end{bmatrix}. \tag{2}$$

Here $B$ represents a linear transform. Note after PCA preprocessing, different components $s_i + e_i$ or $z + e_z$ are still independent and at most only one of the factors is Gaussian. As a result, the ICA identifiability theorem [2] can be directly applied on $\boldsymbol{x}$ to unmix the independent components, which means the confounding factors are identifiable, up to a permutation:

$$W^{\text{ICA}} \hat{\boldsymbol{x}} \overset{\text{i.i.d}}{\sim} \begin{bmatrix} s_1 + e_1 \\ \vdots \\ s_l + e_l \\ z + e_z \end{bmatrix}, \text{ up to a permutation.} \tag{3}$$

Finally, a statistical test on the difference between untreated and treated distributions for each independent component can be performed to distinguish the confounding factors from the treatment event signal.

Our theoretical justification here reveals that: 1. The (noisy version of) confounder terms are identifiable with the ICA transform up to a permutation; 2. The contributions of noise terms are not distorted as the relative ratio between $s, z$ and $e$ are preserved in the output. These two points supports the validity of using CINEMA-OT for matching according to the identified (noisy) confounders.

Moreover, we are able to show that in the same setting, a non-linear neural network based architecture named conditional (variational) autoencoder (used as the key component in scGen [3], compositional autoencoder [4], and contrastiveVI [5] along with other tools) can fail to identify the confounder signals.

In the model of conditional (variational) autoencoders, the latent space can be represented as two parts $\boldsymbol{s} = [\boldsymbol{s}_0; z]$. First is the basal embedding $\boldsymbol{s}_0$, where the embedding of cell distributions from different conditions overlap; The other is the treatment signal $z$, which can be transformed into a treatment associated embedding. The model can be written as the following form:

$$\boldsymbol{x} \sim \int P_\theta(\boldsymbol{x}|\boldsymbol{s}_0, z)P(\boldsymbol{s})d\boldsymbol{s} \tag{4}$$

Here the setting of Gaussian $\boldsymbol{s}$ corresponds to a variational autoencoder design and a uniform $\boldsymbol{s}$ over the whole space corresponds to the vanilla autoencoder design. In our context, we suppose the conditional (variational) autoencoder is optimized by the reconstruction loss with respect to $\hat{\boldsymbol{x}}$ in eq (2) with / without Gaussian constraint:

$$(\hat{\theta}, \hat{\boldsymbol{s}}) = \text{argmin}_{\theta,\boldsymbol{s}}||F_\theta(\boldsymbol{s}_0, z) - \hat{\boldsymbol{x}}|| \tag{5}$$

Even we assume the function above is perfectly optimized, and the learned $\boldsymbol{s}_0$ is indeed independent of $z$, $\boldsymbol{s}_0$ can be still an arbitrary transform of the noisy confounder signals. This is based on the following fundamental result from [6, 7]:

**Theorem.** *[6, 7] Let $\boldsymbol{x}$ be a d-dimensional random vector of any distribution. Then there exists a transformation $F : \mathbb{R}^d \to \mathbb{R}^d$ such that the components of $\boldsymbol{x}' := F(\boldsymbol{x})$ are independent, and each component has a standardized Gaussian distribution. In particular, $\boldsymbol{x}'_1$ equals a monotonic transformation of $\boldsymbol{x}_1$.*

The above theorem means, for any arbitrary injective transformation $F([s_1 + e_1, ..., s_l + e_l]^T)$, its first component can be used as the first independent component in $\boldsymbol{s}_0$. This immediately leads to two observations: 1. The confounder terms are no longer identifiable up to a permutation; 2. Even though the information of confounder can be preserved up to a injective transformation, the distance measure on the latent space can be dramatically distorted. To see why the second point holds, suppose $s_1$ is a binary r.v. with $P(s_1 = 1) = P(s_1 = -1) = 0.5$, and $\frac{Var(s_1)}{Var(e_1)}$ is a sufficiently large constant. We can see in this case the distance based on $s_1 + e_1$ is almost determined by $s_1$; however, the distance measure after an injective transform can be almost determined by $e_1$ (Supplementary Figure 1). In summary, our theoretical analysis suggests that in the model setting, the classical ICA can give consistent distance measures based on confounder space, while the non-linear conditional autoencoder approaches suffer from non-identifiability and distance distortion, leading to less meaningful latent spaces.

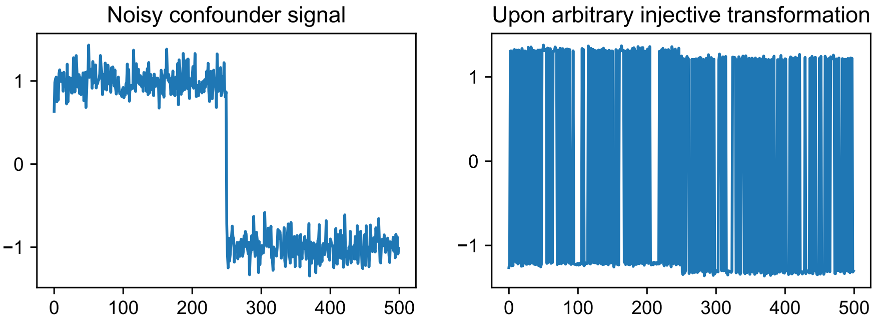


Supplementary Figure 1: An example of distance distortion with noisy confounder signals.

Empirically, we also observe our method can successfully reveal confounding variations even with non-linear interactions between confounding factors and treatment events. In this case, the causal matching may be performed according to a non-one-to-one transform of the full confounder signal, which is not consistent with the full confounder distribution but still meaningful in preserving a part of confounder information. More specifically, it interpolates between a single-cell level causal matching and a cluster/population-level causal matching.

Finally, given the identified confounding factors, the problem of treatment effect estimation can be solved by standard potential outcome framework. The framework has mainly four assumptions, which are discussed in detail in causal inference textbooks [8]:

1. Stable Unit Treatment Assumption (SUTVA): Samples are independent without interference; 2. Ignorability: there are no unmeasured confounders; 3. Consistency; 4. Positivity: for a given confounder, the probability of perturbation is neither 0 or 1.

#### References


[1] Dan Shen, Haipeng Shen, and J Marron. A general framework for consistency of principal component analysis. *Journal of Machine Learning Research*, 2016.

[2] Pierre Comon. Independent component analysis, a new concept? *Signal processing*, 36(3): 287–314, 1994.

[3] Mohammad Lotfollahi, F Alexander Wolf, and Fabian J Theis. scgen predicts single-cell perturbation responses. *Nature methods*, 16(8):715–721, 2019.

[4] Mohammad Lotfollahi, Anna Klimovskaia Susmelj, Carlo De Donno, Yuge Ji, Ignacio L Ibarra, F Alexander Wolf, Nafissa Yakubova, Fabian J Theis, and David Lopez-Paz. Compositional perturbation autoencoder for single-cell response modeling. *BioRxiv*, 2021.

[5] Ethan Weinberger, Chris Lin, and Su-In Lee. Isolating salient variations of interest in single-cell transcriptomic data with contrastivevi. *bioRxiv*, 2021.

[6] Aapo Hyvärinen and Petteri Pajunen. Nonlinear independent component analysis: Existence and uniqueness results. *Neural networks*, 12(3):429–439, 1999.

[7] Ilyes Khemakhem, Diederik Kingma, Ricardo Monti, and Aapo Hyvarinen. Variational autoencoders and nonlinear ica: A unifying framework. In *International Conference on Artificial Intelligence and Statistics*, pages 2207–2217. PMLR, 2020.

[8] Guido W Imbens and Donald B Rubin. *Causal inference in statistics, social, and biomedical sciences*. Cambridge University Press, 2015.

### 2 Supplementary figures

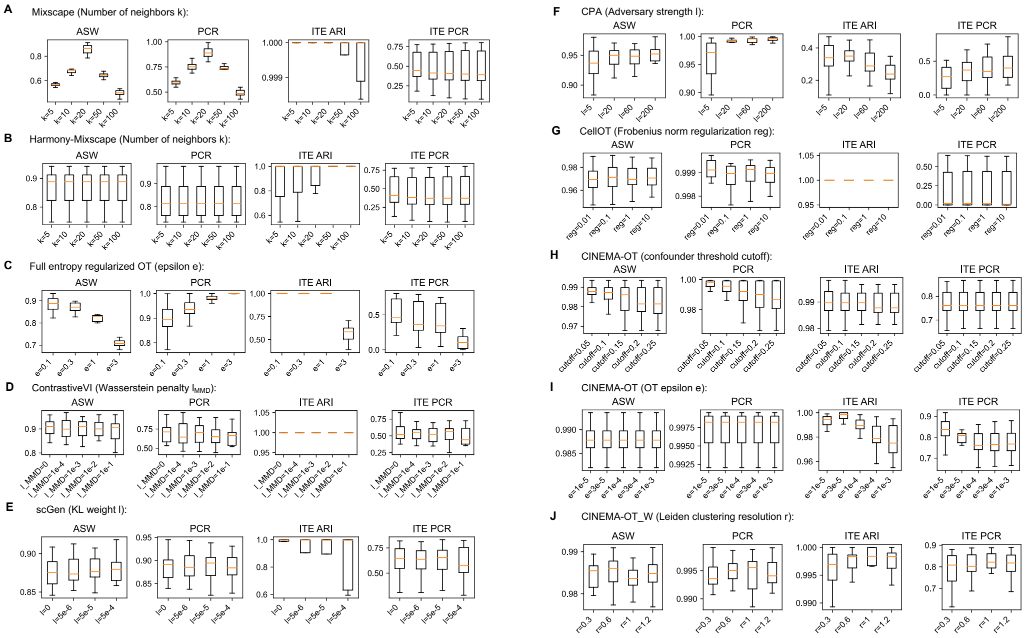


Supplementary Figure 2: Parameter sweep analysis for different single-cell level treatment effect analysis methods.

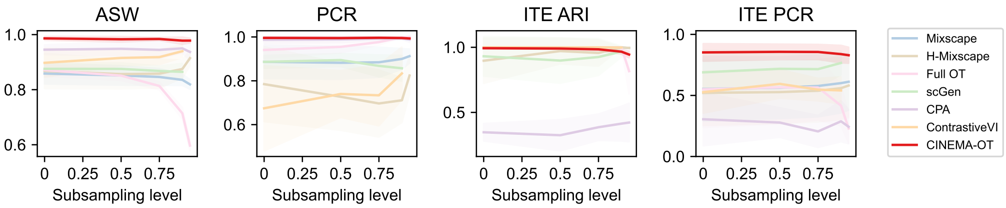


Supplementary Figure 3: Comparison for different single-cell level treatment effect analysis methods' performance as the sparsity level of the dataset increases. Here the result of scGen and ContrastiveVI in the setting of subsampling level = 0.95 is not shown, as they fail to converge or return NaN in some datasets.

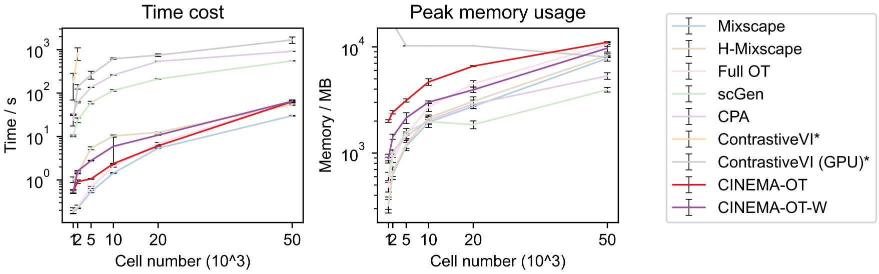


Supplementary Figure 4: Running time and peak memory usage comparison for different single-cell level treatment effect analysis methods. *: These methods are tested on high performance clusters, therefore the performance may not be directly comparable.

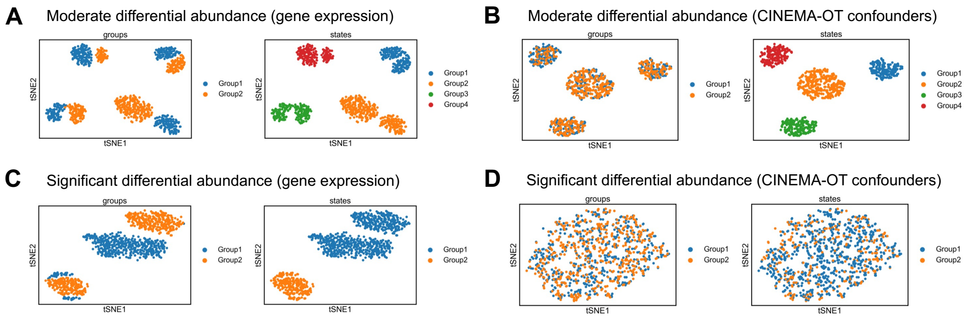


Supplementary Figure 5: CINEMA-OT method still identifies correct confounder in data with moderate differential abundance but fails in data with significant differential abundance.

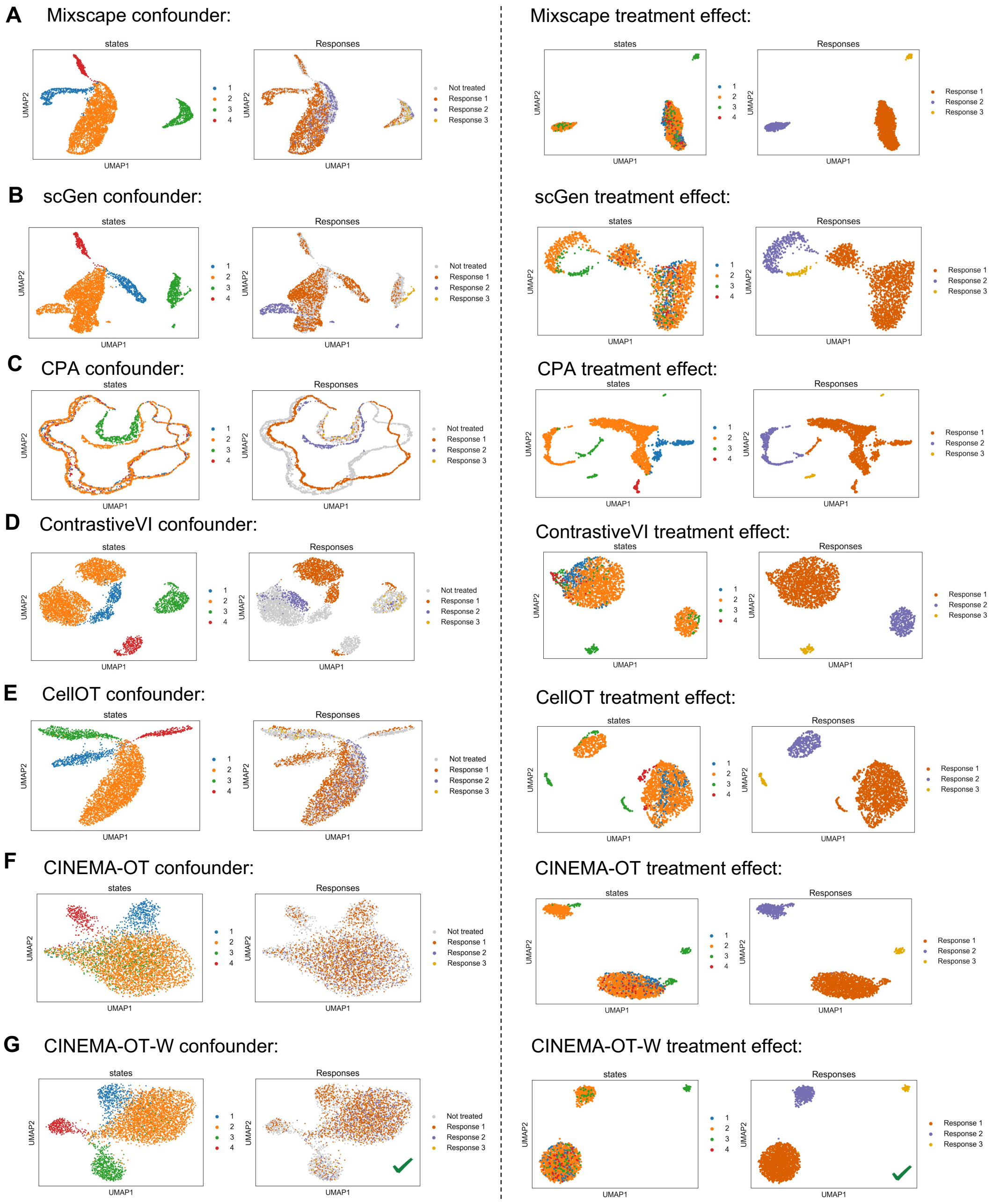


Supplementary Figure 6: CINEMA-OT validation on a simulated dataset example with differential abundance. **A-G.** UMAP visualizations of confounder space (left) and treatment effect space (right) in an simulated dataset with differential abundance ratio = 0.25.

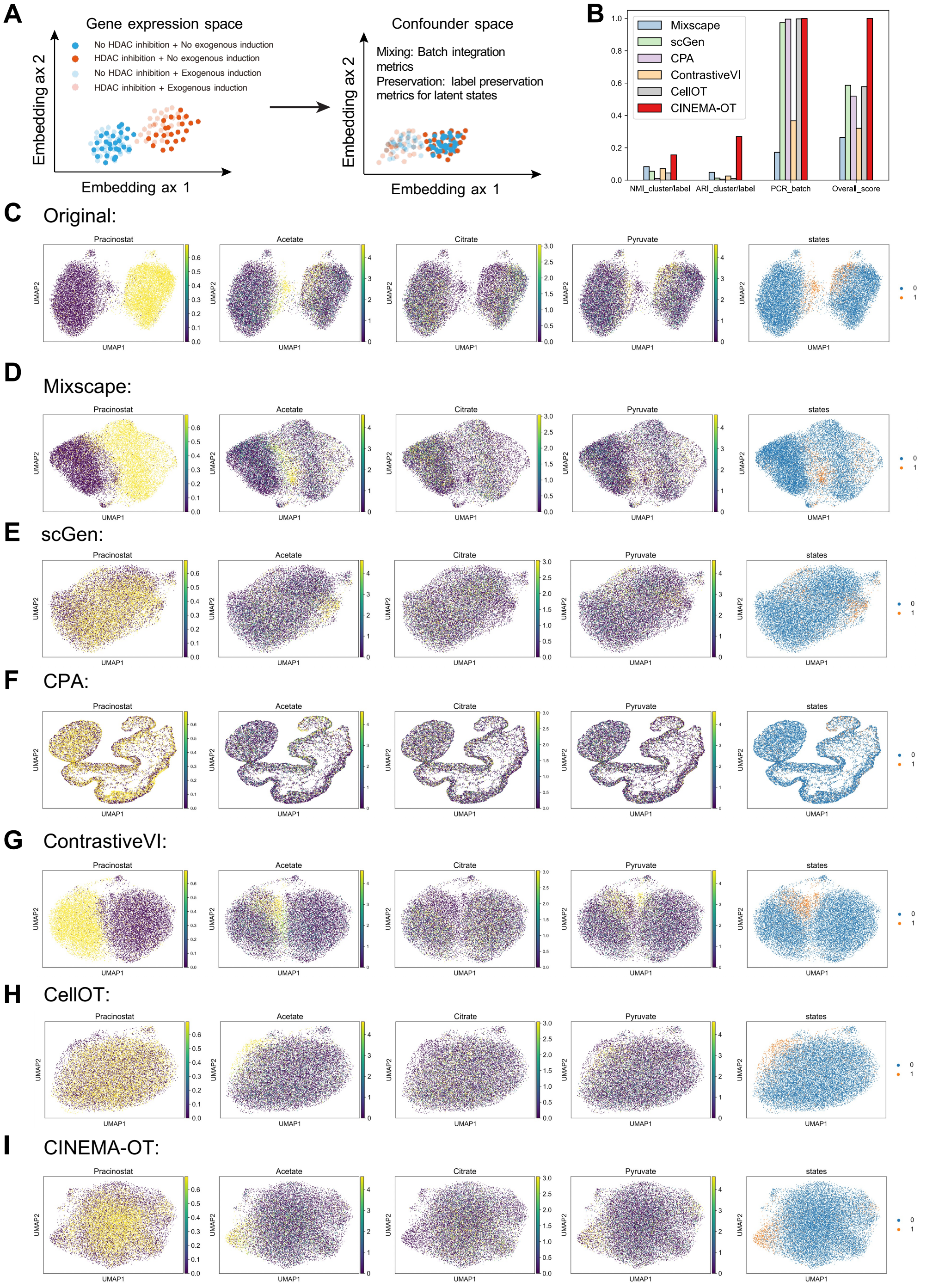


Supplementary Figure 7: CINEMA-OT validation on Sciplex data. **A.** UMAP visualizations of different covariates (Cell type, donor batch, and AD/control) and SPP1 expression. **B-I.** Different methods' confounder space visualization and treatment effect visualization.

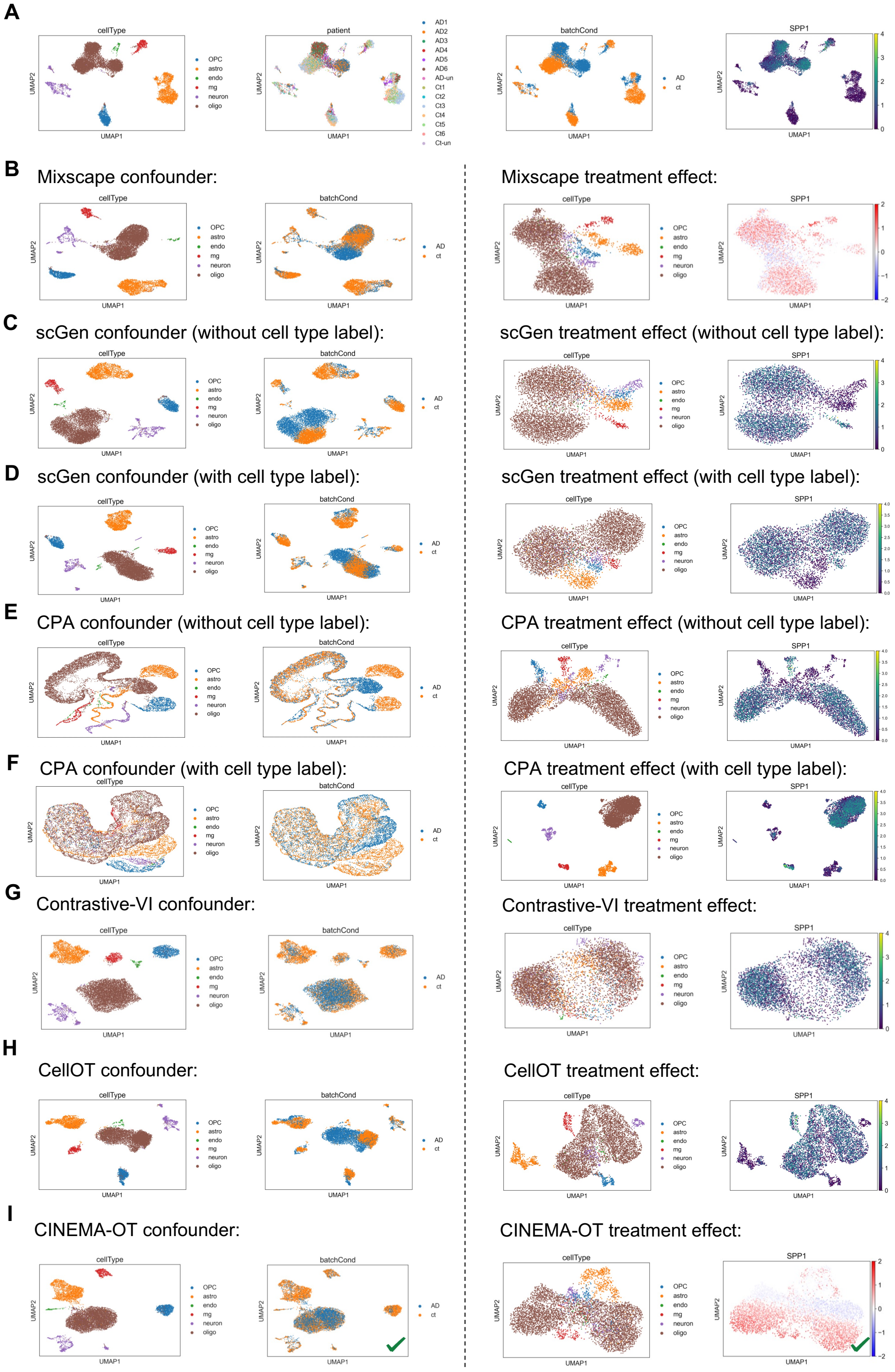


Supplementary Figure 8: CINEMA-OT validation on Alzheimer's data. **A.** UMAP visualizations of different covariates (Cell type, donor batch, and AD/control) and SPP1 expression. **B-I.** Different methods' confounder space visualization and treatment effect visualization.

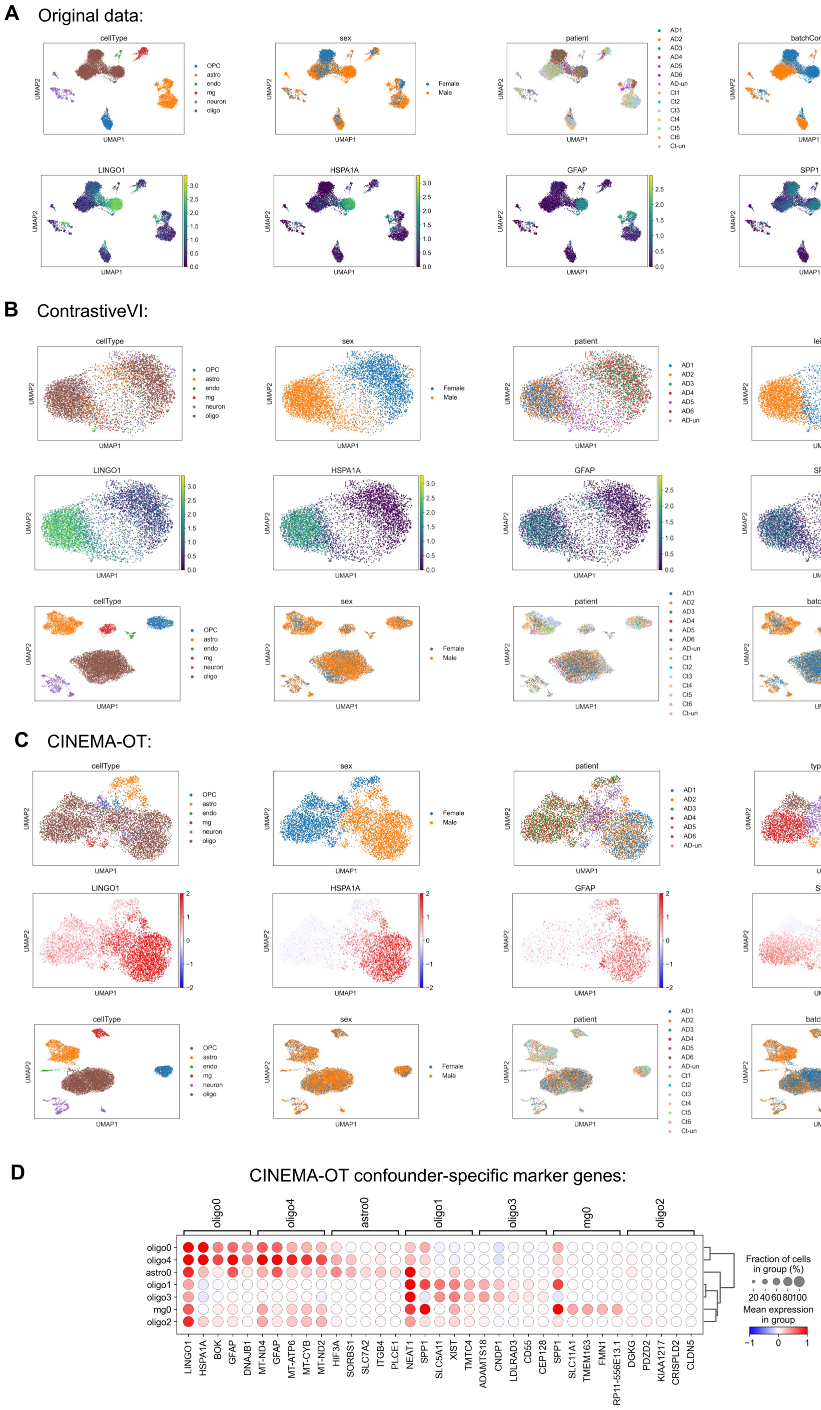


Supplementary Figure 9: Additional comparisons of ContrastiveVI and CINEMA-OT in the Alzheimer scRNA-seq dataset. **A.** UMAP visualizations of different covariates (Cell type, donor sex, donor batch, and AD/control) and expressions of different cell-type specific AD marker genes. **B.** ContrastiveVI response space visualization for covariates and leiden subclusters (first line), state-specific gene visualization in the response space (second line), and confounder space visualization for covariates (third line). **C.** CINEMA-OT response space visualization for covariates and Leiden subclusters (first line), state-specific gene visualization in the response space (second line), and confounder space visualization for covariates (third line). **D.** Response-cluster specific genes identified by CINEMA-OT.

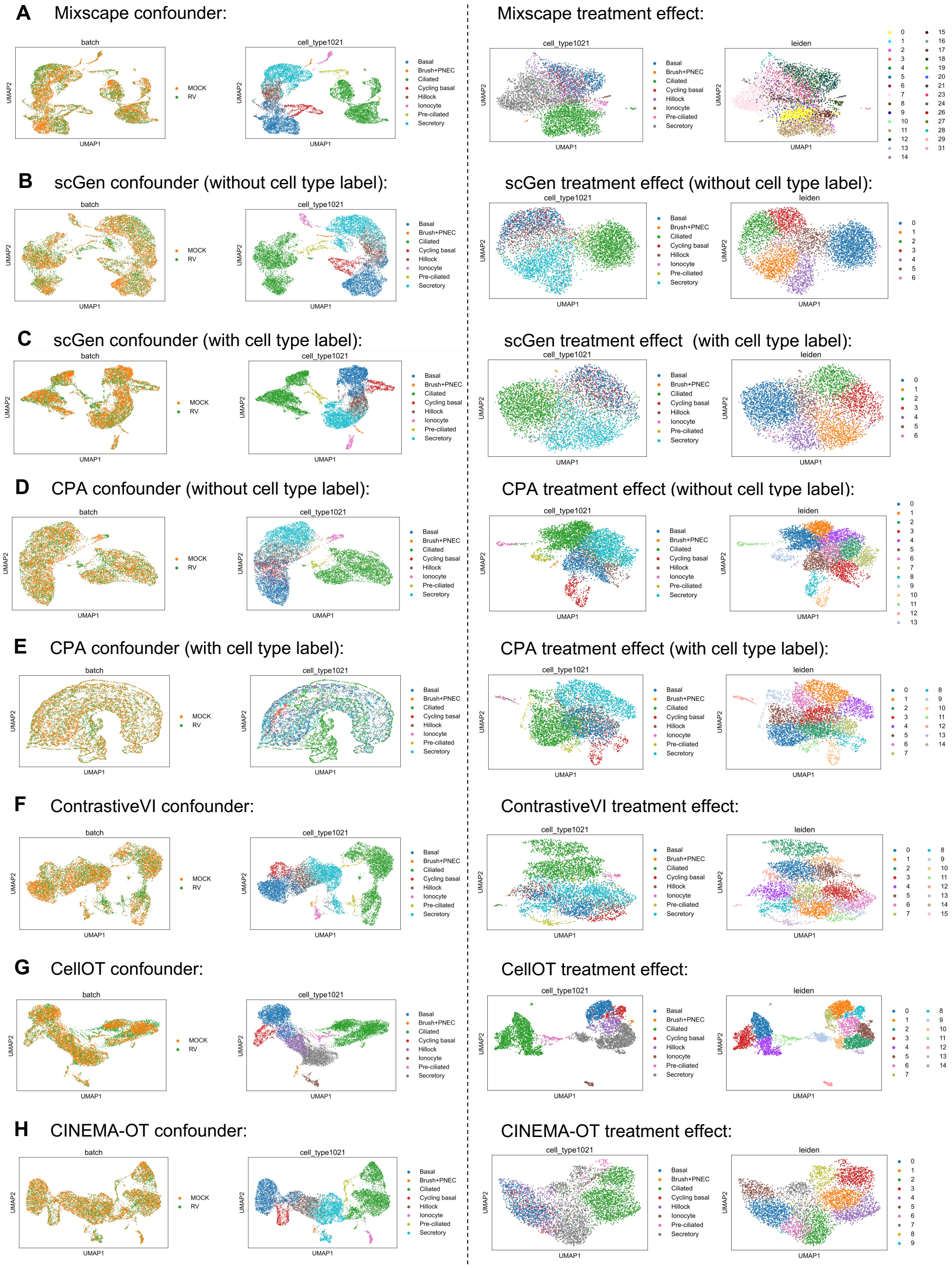


Supplementary Figure 10: CINEMA-OT validation on the Rhinovirus infection data estimating the causal effect of RV. **A-H.** Different methods' confounder space visualization and treatment effect visualization.

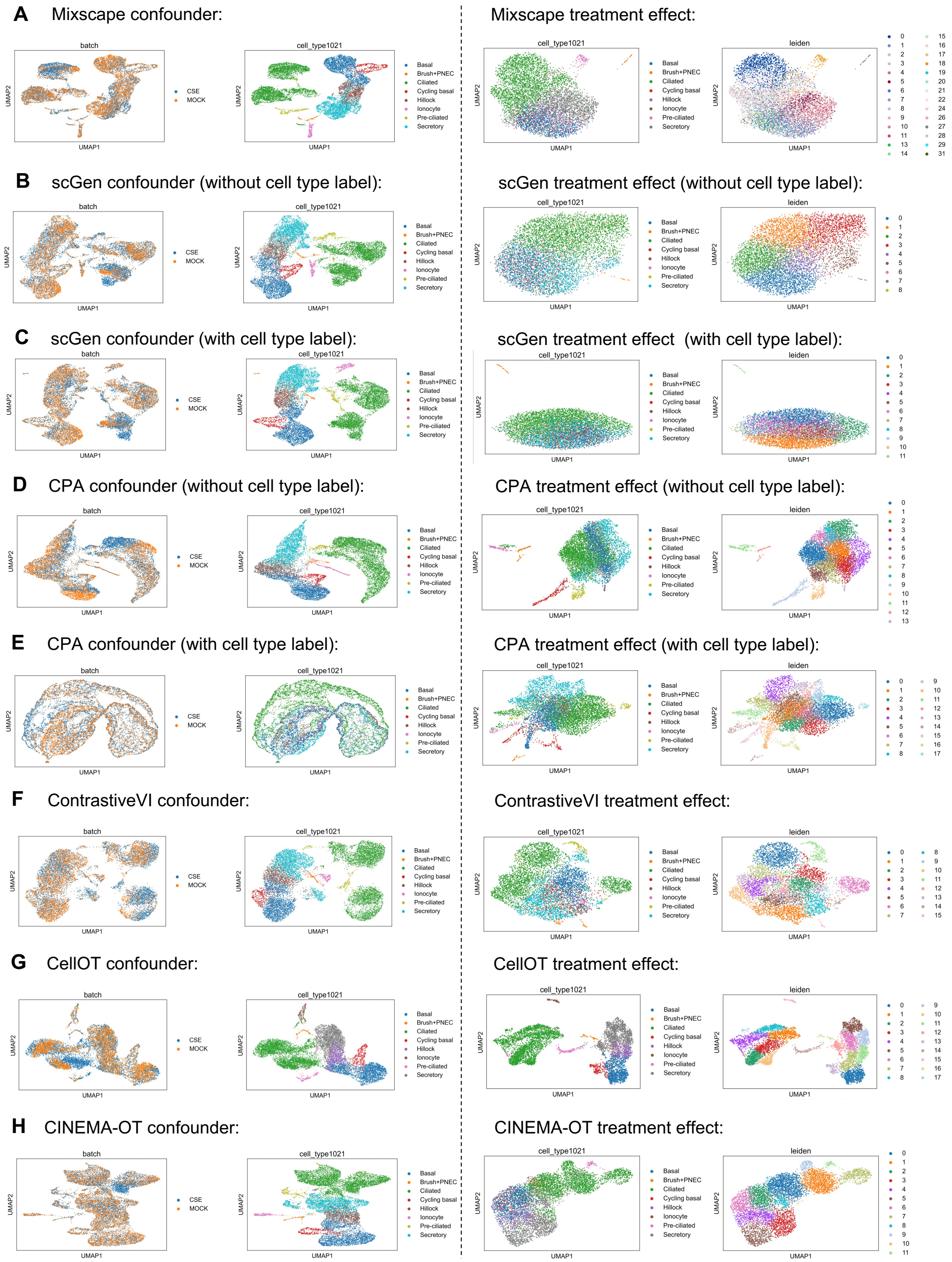


Supplementary Figure 11: CINEMA-OT validation on the Rhinovirus infection data estimating the causal effect of CSE. **A-H.** Different methods' confounder space visualization and treatment effect visualization.

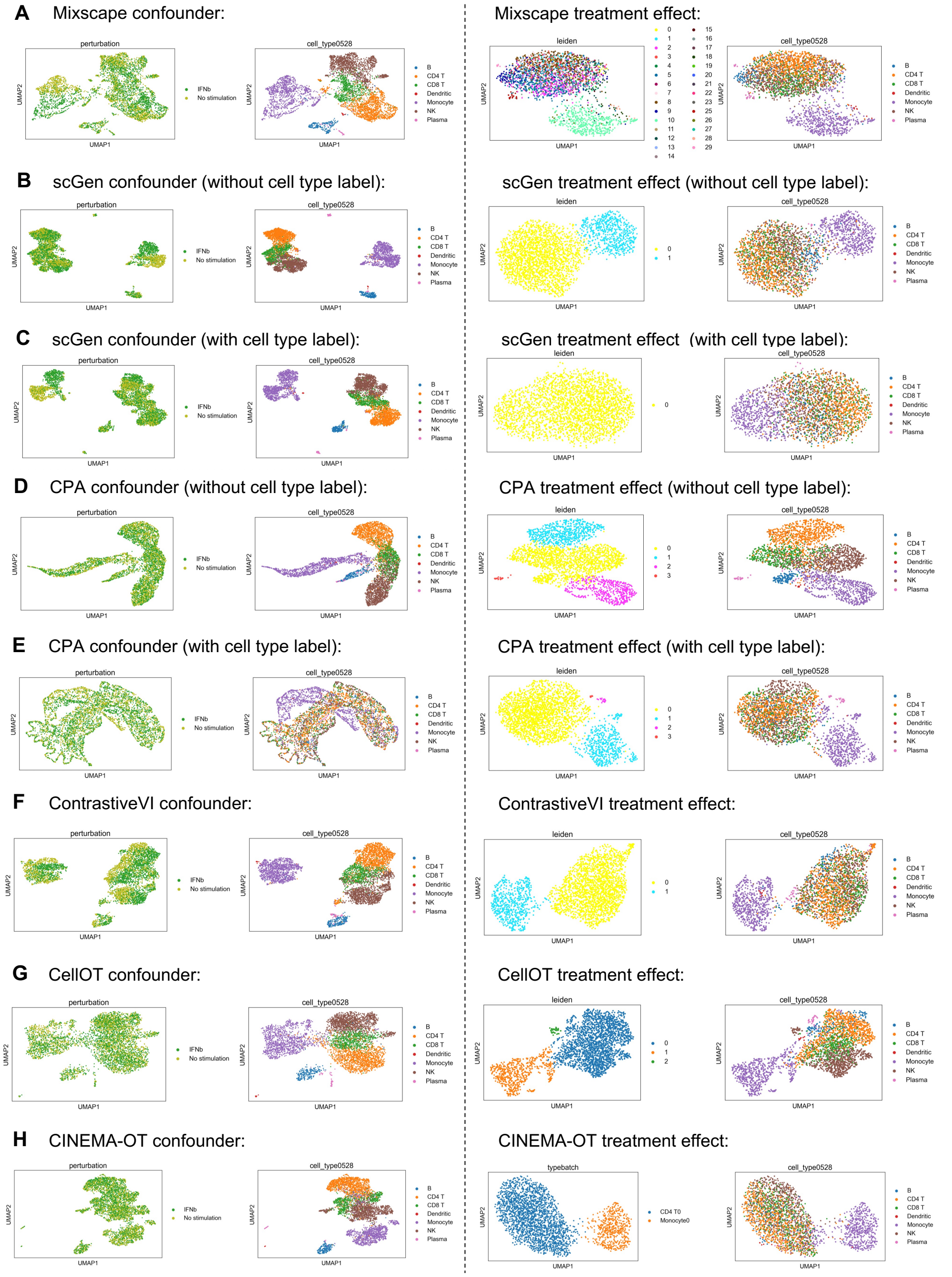


Supplementary Figure 12: CINEMA-OT validation on the interferon data. **A-H.** Different methods' confounder space visualization and treatment effect visualization.

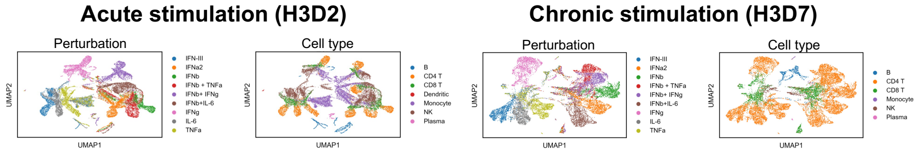


Supplementary Figure 13: UMAP visualizations of batch-wise CINEMA-OT counterfactual space, colored by perturbation and cell type.

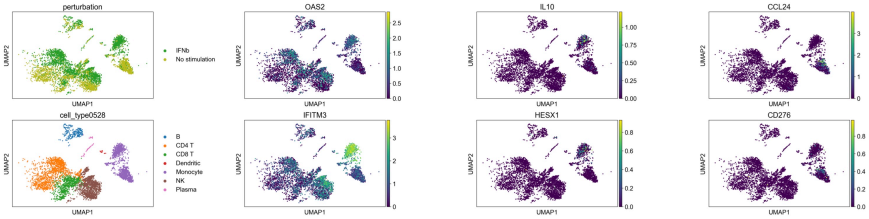


Supplementary Figure 14: Visualizations of identified genes with significant confounder-dependent treatment effect, following Figure 5F.

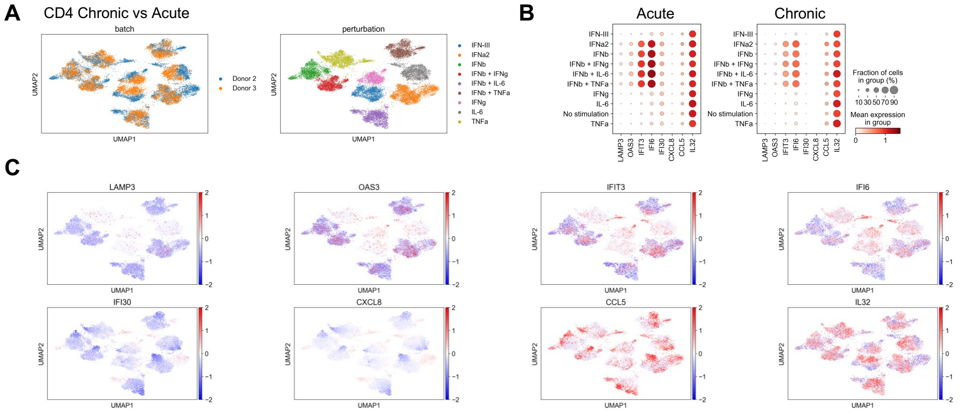


Supplementary Figure 15: Comparison of CD4 T cells gene expression change between chronic stimulation condition and acute stimulation condition across different conditions. **A.** UMAP visualizations of CD4 T cell response space pooled over different perturbations. **B.** Dot plots of selected genes identified by CINEMA-OT. **C.** Single-cell expression differences of selected genes visualized in the UMAP space.

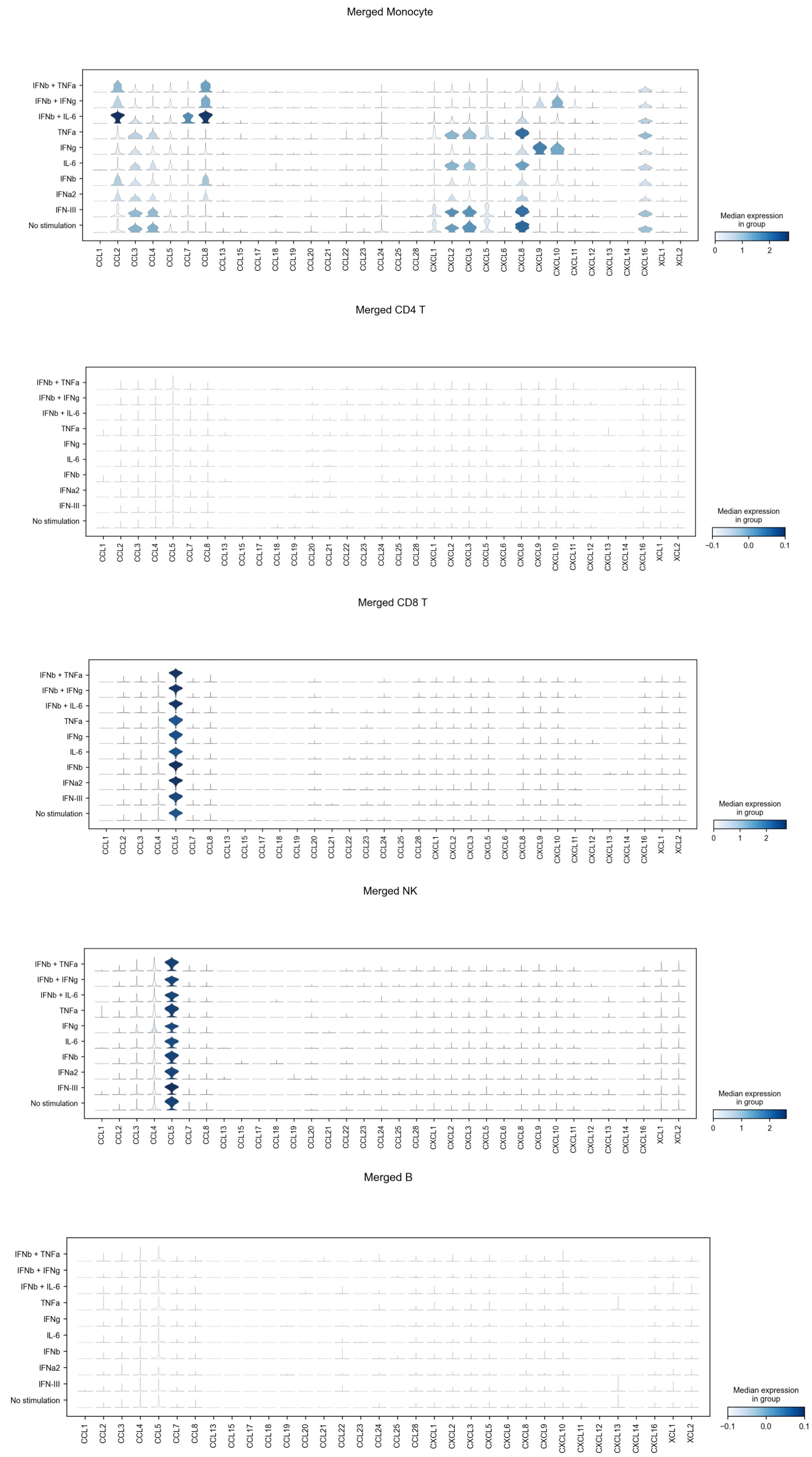


Supplementary Figure 16: Comparison of different different cell types' chemokine response. Systematic differential expressions of chemokines across interferon perturbations are only observed in monocytes.
